# Transposable Elements and Homotypic Niches Drive Immune Dynamics and Resistance in Melanoma Epigenetic-based immunotherapy

**DOI:** 10.1101/2025.10.09.679175

**Authors:** Erika Ciervo, Francesco Ceccarelli, Anna Maria Di Giacomo, Piera Grisolia, Alessia Covre, Zein Mersini Besharat, Antonio De Falco, Francesca Pia Caruso, Luigi Laezza, Luigi Ferraro, Gloria Mas Martin, Maria Fortunata Lofiego, Tommaso Sani, Elisabetta Ferretti, Yan Guo, Sean Barry Holden, Roberta Mortarini, Andrea Anichini, Michele Maio, Teresa Maria Rosaria Noviello, Michele Ceccarelli

## Abstract

Melanoma plasticity drives immune evasion and therapy resistance through dynamic cell-state transitions beyond genetic alterations. Epigenetic remodeling critically influences such processes, yet its role in reshaping the tumor ecosystem under therapeutic pressure remains unresolved. Here, we profiled longitudinal biopsies from melanoma patients treated in the phase Ib NIBIT-M4 epi-immunotherapy clinical trial (NCT02608437), testing the combination of a DNMT1 inhibitor with anti-CTLA4 using single-cell multiome and high-resolution spatial transcriptomics. Integrated analyses resolved seven malignant meta-programs, including a rare Wnt/β-catenin–driven melanocytic state and a de-differentiated neural crest–like state enriched in non-responders. Spatial modeling revealed that homotypic clustering stabilizes resistant programs, with neural crest–like cells forming compact, centrally localized niches, whereas Wnt/β-catenin subpopulations displayed a bimodal architecture, either cohesive clusters sustained by adhesion or dispersed, transcriptionally plastic cells. Responders exhibited progressive enrichment of an antigen presentation/interferon program and coordinated remodeling of the tumor microenvironment with T and B cell expansion, whereas tumors from non-responder patients maintained stable composition of neural crest–like clusters. Epigenetic therapy reactivated transposable elements, providing both regulatory signals that prime innate immunity within microenvironment and generating antigens that drive immunoediting and immunogenicity of Antigen presentation/interferon cell states in responders. Finally, NFATC2 emerged as a master regulator of neural crests–like transcriptional phenotypes and promoter of resistance to therapeutic interventions in melanoma patients. NFATC2 perturbation was able to shift tumor cells towards more differentiated and immunogenic states. These findings reveal how epigenetic-based immunotherapy reshapes melanoma ecosystems, provide mechanistic insights into how multiple transcriptional programs promote tumor plasticity and resistance to both combinatorial therapies and immune checkpoint blockade, identify spatial clustering as a principle stabilizing resistant niches, and highlight β-catenin and NFATC2 as actionable vulnerabilities to overcome resistance.

## Introduction

Melanoma is characterized by high heterogeneity and pronounced phenotype switching between cellular states ^1^. This plasticity is driven by nongenetic reprogramming that regulates reversible transcriptional changes ^2,3^, oscillating between multiple differentiation states that resemble different stages of embryonic development ^4^. For example, de-differentiated melanoma cells express programs resembling neural crest embryonic developmental stages associated with resistance to immune therapies ^5,6^. Similarly, tumor-intrinsic programs associated with major histocompatibility complex (MHC) class I downregulation and MITF^low^/AXL^high^ are hallmarks of resistance to immune checkpoint inhibitors (ICIs) and de-differentiated phenotypes ^7^. The mechanisms by which epigenetic reprogramming affects specific melanoma cellular states and how they influence therapy response are poorly understood, also due to the lack of suitable human patient data/clinical data ^8^. Recent studies have demonstrated that reprogramming with epigenetic drugs can alter the immune microenvironment and affect melanoma cancer cells ^9,10^, as well as enhance the response to immune therapies ^11,12^. The main biological activity of epigenetic drugs, such as DNA (cytosine-5)-methyltransferase 1 (DNMT1) inhibitors, is the promotion of antigen gene expression and activation of master factors belonging to innate immunity pathways, including Type I–III interferon (IFN), NFkB, and TLR10, corroborating the notion that the rescue of adaptive immunity by ICI may cooperate with the promotion of innate immunity by the epigenetic reprogramming, thus potentially explaining the clinical activity of the combination therapy ^13^. Using bulk gene expression, methylation, and DNA sequencing, we have recently reported that epigenetic immune reprogramming, achieved by DNMT1 inhibition in combination with immune checkpoint inhibition, promotes the up-regulation of endogenous retroviral elements and IFN-stimulated genes ^12^. We also demonstrated that tumor-intrinsic factors associated with genetic immunoediting, coupled with intra-tumoral immune features, are an efficient predictor of response to both combinatorial therapy and immune checkpoint inhibition. However, bulk analysis lacks cell-type resolution, whereas integrating single-cell technologies would enable dissection of the precise cellular subsets responding to epigenetic and immune-modulatory pressure. Previous studies using single-cell RNA sequencing have revealed distinct transcriptional programs in melanoma cells, including MITF^high^melanocytic and AXL^high^ invasive states, associated with differential drug sensitivity and plasticity ^14^. Neural crest-like and stress-related tumor-specific programs characterize minimal residual disease, highlighting the dynamic nature of melanoma cell state transitions ^15^. Other cellular states associated with antigen-presenting and undifferentiated phenotypes were also uncovered with single-cell technologies ^1^. Pozniak et al. ^16^ recently integrated multimodal single-cell and spatial transcriptomic data from melanoma patients treated with immune checkpoint inhibitors, revealing how transitions between melanoma cell states, particularly the shift from melanocytic to mesenchymal-like phenotypes, are associated with immune evasion and therapeutic resistance. The lack of genetic alterations supporting the recurrent melanoma cellular states observed in these studies highlights the role of nongenetic mechanisms in influencing phenotypic switching between tumor cellular states in melanoma, as well as their cross-talk with the microenvironment and association with response to treatment. To uncover rare or transitional cellular phenotypes, lineage-specific transcriptional rewiring, and spatially confined tumor-immune network, ultimately refining the evolution of cellular states under immune therapy and elucidating resistance mechanisms not apparent at the bulk level, we have generated single-cell multiome coupled with high-resolution spatial transcriptomics of longitudinal biopsies from melanoma patients treated in the Phase Ib NIBIT-M4 trial, with the association of the anti-CTLA4 ipilimumab, and the DNMT1 guadecitabine ^11,12^. Here, we show how epi-immunotherapy reshapes melanoma ecosystems. We demonstrate that clinical response/benefit is associated with progressive enrichment of an Antigen presentation/Interferon program, whereas resistance is marked by the persistence of a Neural crest–like state. Consistent with recent evidence that homotypic clustering stabilizes glioblastoma states and constrains transcriptional heterogeneity ^17^, we observed that resistant neural crest–like programs form compact homotypic niches enriched in non-responders. Our data extend this principle by showing that different melanoma states adopt distinct spatial strategies: while Neural crest-like programs rely on cohesive niches for stability, Wnt/β-catenin cells can exist in both clustered and dispersed configurations, with dispersion linked to higher plasticity. Moreover, we show that transposable element (TE) reactivation driven by epigenetic-based immunotherapy contributes both to regulatory and antigenic signals, highlighting a dual role in shaping tumor–immune dynamics. Finally, we identify NFATC2-driven plasticity as a central therapeutic vulnerability, whose inhibition may promote re-differentiation and overcome the resistant Neural crest–like state.

## Results

### Longitudinal single-cell multiome profiling of the NIBIT-M4 clinical trial

To map the dynamic evolution of distinct malignant cell subpopulations and the tumor microenvironment (TME) over the course of epigenetic therapy in melanoma, we performed longitudinal profiling using the 10x Genomics Chromium Single Cell Multiome platform on tumor biopsies collected at baseline (week 0) and during treatment (weeks 4 and 12) from melanoma patients treated in the NIBIT-M4 trial (**Figure 1a**). We generated paired single-nucleus RNA-seq (snRNA-seq) and ATAC-seq (snATAC-seq) libraries across thirteen tumor specimens obtained from five patients, two responder (R) and three non-responder (NR) patients. Following integrated quality control across both modalities, we retained 38,029 high-quality cells with matched transcriptomic and chromatin accessibility profiles, comprising 58,670 cells that passed RNA filters and 55,850 cells that passed ATAC filters.

**Figure 1.**
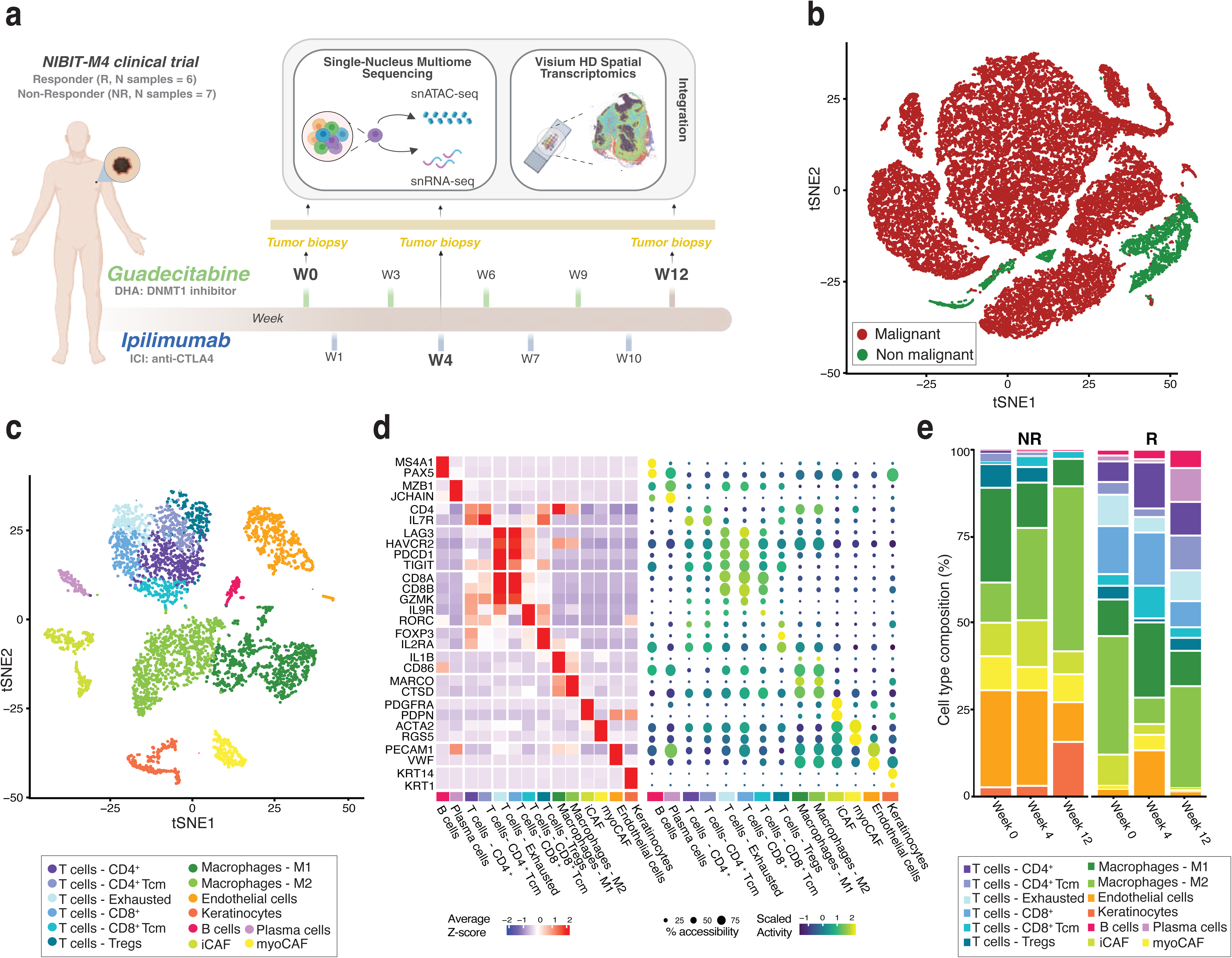
Multiomic characterization of the NIBIT-M4 cohort. **a.** Schematic overview of the study design. Tumor biopsies were collected from patients classified as Responders (R, n = 6) and Non-Responders (NR, n = 7) at baseline (W0), week 4 (W4), and week 12 (W12), and profiled using single-nucleus Multiome sequencing (snRNA-seq and snATAC-seq) along with Visium HD Spatial Transcriptomics. **b.** t-distributed stochastic neighbor embedding (t-SNE) of multi-modal snRNA-seq and snATAC-seq data colored by malignant and non-malignant cell populations. **c.** t-SNE of the tumor microenvironment (TME) using snRNA-seq data, colored by cell types. Abbreviations: Tcm, T central memory; Tregs, regulatory T cells; iCAF, inflammatory cancer-associated fibroblasts; myoCAF, myofibroblastic cancer-associated fibroblasts. **d.** Expression and activity of marker genes across cell types shown as average z-score expression (left) and scaled gene activity (dot plot, right) **e.** Bar plot showing TME cell type composition in NR and R patients across time points.

Cells were classified as malignant (n = 34,822) based on inferred copy number alterations (CNAs) from gene expression data ^18^ (**Supplementary Figure 1a**). Unsupervised dimensionality reduction, integrating both transcriptomic and chromatin accessibility modalities, consistently separated non-malignant from malignant compartments, as visualized using t-distributed Stochastic Neighbor Embedding (t-SNE) (**Figure 1b**), malignant cells were largely intermixed between time points, indicating that global transcriptional programs were conserved across the study period. Nonetheless, specific clusters appeared enriched for cells collected at week 4, reflecting the abundance of cells collected at week 4 (**Supplementary Figure 1b**).

To characterize changes in the TME during treatment, non-malignant cell types (n = 4,968) were identified through clustering analysis followed by cell type annotation (**Figure 1c** and **Supplementary Table 1**). Given the limited availability of cell-type-specific chromatin accessibility markers, annotation was performed using the snRNA-seq modality, leveraging validated reference gene signatures and canonical marker gene expression (**Figure 1d, left**). We identified multiple T cell subpopulations, including CD8^+^, CD4^+^, exhausted, regulatory (Tregs), and central memory (Tcm) T cells, as well as B cells and plasma cells. Macrophages were further classified into M1 and M2 polarized states, while the stromal compartment comprised cancer-associated fibroblasts (CAFs), distinguished into inflammatory CAFs (iCAFs) and myofibroblastic CAFs (myoCAFs)^19,20^. Additionally, endothelial cells and keratinocytes were also detected. The snATAC-seq data corroborated the reliability of TME cell type annotation derived from the transcriptome data. The same marker genes that exhibited cell type-specific expression in the RNA modality also showed consistent patterns of chromatin accessibility in the corresponding TME clusters, as measured by gene accessibility scores (**Figure 1d, right**).

As a longitudinal/prospective clinical trial, this study enabled the longitudinal tracking of temporal dynamics within the tumor ecosystem at single-cell resolution. Combination therapy induced a progressive immune engagement (**Figure 1e**) and functional remodeling of CD4^+^ and CD8^+^ T cells, as indicated by the transcriptional and epigenetic activation of the PD-1 signaling pathway (**Supplementary Figure 1c**), indicative of a sustained and coordinated immune engagement over time and consistent with previous findings from bulk analyses ^12^. We further observed that the composition of TME cell populations varied over time within response groups (**Figure 1e**). In NR patients, the TME remained largely immunosuppressive throughout treatment, characterized by a dominance of M2-polarized macrophages, limited T cell infiltration, and persistent stromal elements such as iCAFs and keratinocytes, reflecting poor immune activation. In contrast, R patients exhibited dynamic TME remodeling. By week 4, there was a notable increase in CD8^+^ and CD4^+^ T cells, exhausted T cells, and M1 macrophages, suggesting early signs of immune activation. This trend further intensified by week 12, with a marked expansion of T cells accompanied by the emergence of B cells and plasma cells, consistent with a coordinated and sustained anti-tumor immune response. Consistently, expression levels of key immune checkpoint genes were significantly elevated in T cells from R patients over time (**Supplementary Figure 1d),** reinforcing the sustained immune activation and adaptive feedback typical of effective immune responses.

Overall, our longitudinal single-cell analysis confirms that effective response to epi-immunotherapy in melanoma, combining DNMT1 inhibition with ICI, is associated with dynamic and coordinated remodeling of the TME, characterized by progressive immune activation and diversification over the course of treatment.

### Dissecting the heterogeneity of melanoma cell states under epi-immunotherapy

To resolve the complexity of melanoma under epi-immunotherapy, we applied a non-negative matrix factorization (NMF) framework to snRNA-seq data from malignant cells, enabling the identification of shared transcriptional programs ^21^. This analysis revealed seven consensus meta-programs (MPs), representing distinct cellular identities or activity states that were conserved across tumors. Each of the seven NMF-derived MPs exhibited unique gene expression patterns, as well as strong internal correlation and gene overlaps (**Figure 2a** and **Supplementary Figure 2a**). Subsequently, we assigned each malignant cell to a specific MP based on the highest MP signature score (see *Methods*) (**Supplementary Figure 2b)**. Cells falling below a simplicity score threshold of 0.1, and therefore not confidently classified, were deemed undefined and excluded from subsequent analyses.

**Figure 2.**
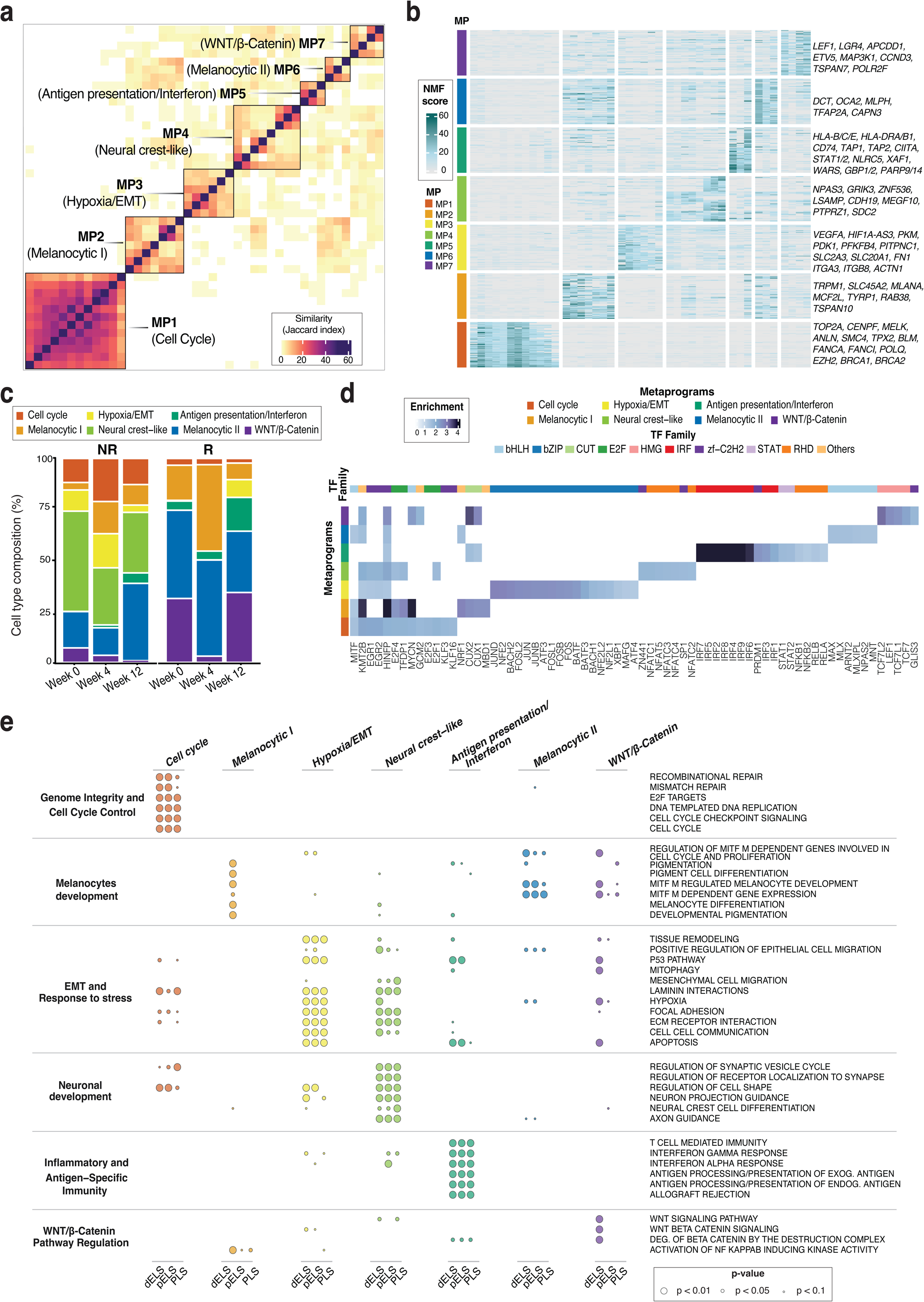
Characterization of metaprograms and their regulatory activity. **a.** Similarity matrix of NMF programs based on gene overlap (Jaccard index). Programs are clustered and grouped into metaprograms (MPs), each annotated with a functional description derived from the top 50 genes. **b.** Heatmap showing NMF scores of genes within MP signatures. Rows represent genes and columns represent programs composing each MP. Selected genes for each MP are highlighted. **c.** Bar plot of MP composition in NR and R patients across time points. **d.** Motif enrichment map of TF binding motifs (columns) across MPs (rows), highlighting candidate regulators of transcriptional states. TFs are color-coded according to their families. **e.** Bubble plot of pathway enrichment for MP-specific TF target genes identified through cCRE–TF–gene linkage analysis. Bubble size indicates enrichment significance (*p*-value), and color denotes the corresponding MP.

Taking advantage of the joint multi-omic resolution of our tumor specimens under epigenetic therapy, we first assessed the extent to which transcriptional states from snRNA-seq are reflected in chromatin accessibility profiles from snATAC-seq, quantified as the percentage of cells with correctly predicted annotations via label transfer across the two modalities (**Supplementary Figure 2c**). The extent to which accessibility profiles predict transcriptional states can serve as a measure of epigenetic plasticity, where cells with uncertain predictions are indicative of more plastic states ^22,23^. Malignant cells exhibited greater prediction variability than non-malignant cells in the TME, with chromatin accessibility often compatible with more than one transcriptional state. The degree of such flexibility varied across tumor cell states: while some MPs retained stable and concordant identities across modalities, others displayed higher heterogeneity. In particular, MP1 was characterized by the highest plasticity. These findings are in line with recent work in other tumor types ^23^, supporting the notion that tumor cell states span a continuum shaped by chromatin remodeling, transitioning between states.

Then, to interpret the biological relevance of each MP, we analyzed the top 50 genes ranked by NMF scores for each MP (**Figure 2b** and **Supplementary Table 2**) and assessed their functional enrichment to elucidate underlying biological processes (**Supplementary Table 3**). When interpretation was challenging, we also analyzed the differential expression profile of each MP compared to the others (**Supplementary Table 4**), followed by functional enrichment analysis (see *Methods,* **Supplementary Table 5**).

MP1 was characterized by genes involved in cell proliferation and DNA repair mechanisms, highlighting biological processes such as cell cycle regulation, chromosomal stability, DNA damage response, and mitosis. It expresses key genes such as *TOP2A*, *MELK*, and *ANLN,* which regulate mitotic progression, spindle assembly, and cytokinesis, hallmarks of rapidly dividing melanoma cells ^24,25,26^. The enrichment of DNA repair factors, including *BRCA1*, *BRCA2*, *FANCA*, *FANCI*, *POLQ*, and *BLM* suggests that melanoma cells in this program rely heavily on homologous recombination and replication stress response pathways to preserve genomic integrity under proliferative stress ^27^. This metaprogram, hereafter referred to as *Cell Cycle*, recapitulates a proliferative transcriptional phenotype, consistently identified in human tumor biopsies, including melanoma ^4,14,16,21^ (**Supplementary Figure 2d**).

MP2, hereafter referred to as *Melanocytic I*, was defined by a melanocytic-like differentiated phenotype, characterized by the expression of genes such as *TRPM1*, *SLC45A2*, *MLANA*, *TYRP1*, and *RAB38*, which are involved in melanin biosynthesis, vesicle trafficking, and melanocyte differentiation ^28–31^. Furthermore, pathways regulated by *MITF* were significantly enriched among the top-ranked gene sets (**Supplementary Table 3**), underscoring the central role of MITF-driven transcriptional programs in defining this metaprogram. This is in line with the established melanocytic differentiation axis associated with MITF activity ^4,14,16,21^ (**Supplementary Figure 2d**).

MP3 was associated with cellular responses to hypoxia and stress, and was thus designated as *Hypoxia/EMT*. Key genes such as *VEGFA*, *HIF1A-AS3*, and *PKM* facilitate angiogenesis and regulate hypoxia-inducible factor (HIF) signaling, enabling adaptation to low oxygen levels ^32–34^. Moreover, a metabolic stress phenotype was evident through the expression of *PDK1*, *PFKFB4*, *PITPNC1, SLC2A3*, and *SLC20A1* ^35,36,37,38^. Additionally, genes such as *ITGA3*, *ITGB8*, *FN1*, and *ACTN1* mediate cell adhesion and extracellular matrix (ECM) remodeling ^39^, further supporting the invasive and hypoxia-adapted phenotype of these melanoma cells.

MP4 was characterized by the presence of genes involved in neural signaling pathways, including *GRIK3*, a neurotransmission receptor subunit; *ZNF536*, a brain-specific zinc finger protein; *LSAMP*, a neural adhesion molecule; *MEGF10*, a phagocytic receptor highly expressed in glial cells; and *CDH19*, a cadherin contributing to neural adhesion. These features suggest that this metaprogram reflects a de-differentiated state of melanoma cells transitioning toward a neural crest-like phenotype (**Supplementary Figure 2d**), and was therefore named *Neural crest-like*. This MP was enriched exclusively in NR patients (Wilcoxon test, *p* = 0.02: **Figure 2c**), as expected for an invasive phenotype, previously associated with resistance to ICI ^40^. Additionally, the expression of *PTPRZ1* and *SDC2* further supports the acquisition of stem cell-like traits and therapy resistance, potentially mediated by interactions with immunosuppressive cells in the TME. In line with this, cell-cell interaction analysis in NR patients revealed activation of the pro-tumorigenic Pleiotrophin (PTN) signaling pathway ^41^, involving these genes and suggesting a crosstalk between this cell state and CAFs (**Supplementary Figure 2e**). Notably, similar interactions have recently been implicated in resistance to ICI in gastric cancer ^42^.

MP5 exhibits remarkable immune-related traits, with genes involved in the antigen presentation machinery, both MHC I and II, including *HLA-B*, *HLA-C*, *HLA-E*, *HLA-DRA*, *HLA-DRB1, CD74, TAP1, TAP2,* and *CIITA*. Additionally, transcription factors and immune effectors such as *STAT1, STAT2, NLRC5, XAF1, WARS, GBP1, GBP2, PARP9,* and *PARP14*, as well as a suite of interferon-stimulated genes, collectively justifying its designation as the *Antigen Presentation/Interferon* metaprogram. Consistent with previous findings ^16,21^, these tumor cells exhibited an immunologically active phenotype within the TME, contributing to improved response. Accordingly, MP5 was more enriched in R versus NR patients (Wilcoxon test, *p* = 0.07; **Figure 2c**). Cell-cell interaction analysis in R patients further revealed robust activation of both MHC class I and, notably, class II pathways (**Supplementary Figure 2f**). Cells in this program displayed a progressive upregulation of immunoproteasome subunits (*PSMB8, PSMB9, PSMB10*) across treatment time points (**Supplementary Figure 2g**), whereas constitutive proteasome components (*PSMB5, PSMB6, PSMB7*) showed no distinct pattern (data not shown). This suggests that therapy induces an immunoproteasome-dependent antigen processing machinery, which has been previously associated with a favorable ICI response in melanoma ^43^.

MP6 was also associated with a melanocytic-like phenotype, highlighted by the expression of key pigmentation-related and MITF-dependent genes (**Supplementary Table 3**), including *DCT, OCA2, MLPH*, *TFAP2A* and *CAPN3* ^44–49^. A strong correlation and substantial gene overlap further corroborated this phenotypic link with the previously defined *Melanocytic I* (**Figure2b** and **Supplementary Figure 2a**), as well as with other known melanocytic cell states (**Supplementary Figure 2d**). Based on this evidence, we designated MP6 as *Melanocytic II*.

MP7 was marked by strong activation of the Wnt/β-catenin signaling pathway, evidenced by high expression of *LEF1*, a key transcription factor downstream of β-catenin. LEF1 is known to play a central role in melanocytic differentiation, as Wnt/β-catenin signaling is critical for the early specification and maturation of melanocyte precursors from the neural crest ^50,51^. Further support for Wnt pathway activation came from its selective upregulation in MP7 compared to other melanocytic programs (Melanocytic I/II), along with the highest expression of the *CTNNB1* (β-catenin) gene (**Supplementary Figure 2h**). Based on these findings, we classified MP7 as a distinct melanocytic-like metaprogram driven by such signaling, and accordingly named it *Wnt/β-catenin*.

Together, these results reveal a heterogeneous landscape of malignant cell states, each defined by distinct transcriptional and functional traits, that may collectively influence the balance between therapeutic sensitivity and resistance in melanoma under epi-immunotherapy.

### Integrated analysis of epigenetic regulation and transcription factor activity in melanoma cell states under epi-immunotherapy

The availability of paired RNA and ATAC measurements enabled not only the characterization of dynamic transcriptional states and DNA regulatory element activities but also facilitated direct mapping of epigenetic gene regulation to gene expression changes.

To investigate the regulatory mechanisms driving melanoma cell metaprograms and their changes under therapy, we performed transcription factor (TF) motif enrichment analysis on differentially accessible chromatin regions identified from snATAC-seq data. Across metaprograms, we identified an average of 19,787 accessible peaks with significant enrichment (average log_2_FC ≥ 0.58 and adjusted *p-value* < 0.05). To identify key TFs controlling each metaprogram, enriched motifs within these peaks were examined. We prioritized motifs with (i) fold enrichment ≥ 1.3, (ii) retaining the most enriched motif when multiple motifs correspond to the same TF, and (iii) compiling the top 30 most enriched TF motifs per metaprogram. A total of 113 unique TFs were identified across the seven MPs (**Supplementary Table 6**). From these, a subset of 69 well-characterized TFs was visualized (**Figure 2d**) to highlight the transcriptional programs and phenotypic identities of the MPs, as previously defined by snRNA-seq.

*MITF* was confirmed as the principal regulator of the melanocytic MPs, in line with its well-established role in melanocyte differentiation and pigmentation. The Hypoxia/EMT state showed strong enrichment for TFs from the bZIP (basic leucine zipper) family, which are known mediators of stress responses, cellular adaptation, and epithelial-to-mesenchymal transition, as *ATF3* and the FOS/JUN (AP-1) complex and *BACH1* ^52,53^. Similarly, the Antigen Presentation/Interferon metaprogram was marked by specific and strong enrichment of IRF and STAT family TFs, driving an immunologically active phenotype. The Wnt/β-catenin metaprogram featured enrichment of HMG-box family TFs, including *LEF1* and *TCF7L2* ^54^, which are well-known regulators of Wnt signaling (**Supplementary Figure 3a**). Interestingly, the Neural crest-like metaprogram appeared to be predominantly regulated by members of the nuclear factor of activated T cells (NFAT) family, TFs best known for their role in calcium-dependent signaling and involved in the control of T-cell development, and T-cell differentiation ^55^. As a complementary approach, enhancer-driven gene regulatory networks (eGRNs) were reconstructed by integrative analysis of paired snRNA-seq and snATAC-seq multiome data using the SCENIC+ pipeline. This analysis further supported the involvement of the same key TFs identified via motif enrichment in regulating metaprogram-specific gene expression (**Supplementary Figure 3b** and **Supplementary Table 7**).

To further dissect the contribution of key TFs to MP-specific regulatory phenotypes, we conducted an integrative cis-regulatory element (CRE)–TF–gene linkage analysis. For each TF, motif-enriched peaks overlapping ENCODE-annotated promoter- and enhancer-like elements (both proximal and distal) were retained (n = 61,054), with 6.84% mapping to promoter-like signatures (PLS), 17.84% to proximal enhancer-like signatures (pELS), and 75.32% to distal enhancer-like signatures (dELS). Genes linked to these CREs (n = 16,237) were then intersected with MP-specific upregulated genes identified from differentially expression analysis on snRNA-seq data (**Supplementary Table 4**), leading to the identification of a high-confidence set of 1,953 MP-specific target genes of enriched TFs across PLS, pELS, and dELS layers. Downstream pathway analysis on these genes (**Figure 2e** and **Supplementary Table 8**) revealed that melanocyte development-related pathways were enriched across multiple MPs, consistent with the tumor cell origin and largely independent of the CRE category. However, distinct phenotypes emerged when focusing on dELS, particularly for the *Melanocytic I* and *Wnt/β-catenin* MPs, highlighting the potential regulatory importance of distal elements in defining these cell states. *Neural crest-like* cells showed strong enrichment for neural development pathways and EMT-related processes such as focal adhesion and ECM-receptor interaction (*p*-value < 0.01), reinforcing their neural origin and invasive phenotype. The *Antigen Presentation/Interferon* state was highly specific for inflammatory and antigen-specific immunity pathways.

Collectively, these results indicate that distinct TF regulatory networks, mediated via promoter and enhancer elements, underlie the functional heterogeneity and phenotypic specialization of melanoma cell states during epi-immunotherapy.

### Spatial clustering stabilizes resistant tumor states and modulates phenotypic plasticity

To analyze how melanoma cells interact with their microenvironment and how cellular neighborhoods shape tumor heterogeneity and progression, we investigated the heterogeneity and spatial organization of metastatic melanoma lesions under therapy in nine samples from four patients (R n=2, and NR n=2) using 10x Genomics VisiumHD profiling. We performed nuclei segmentation on the high-resolution hematoxylin and eosin (H&E) stained images to delineate individual nuclei (see *Methods*), resulting in an average of 113,366 cells per sample. QC filters were applied to remove cells with high mitochondrial gene content and low complexity; a total of 736,028 cells passed the quality control step, and their gene expression counts were normalized and log_2_-transformed. After dimensionality reduction and graph-based clustering, cells were classified as malignant and non-malignant based on the expression of well-known melanoma markers (*PMEL, TYRP1, DCT, MLANA*). On average, 50% of cells were classified as malignant, and 50% as non-malignant, with fractions of malignant cells and non-malignant cells ranging from 30% to 65% and 35% to 70%, respectively. To uncover the spatial dynamics of melanoma cellular states, we annotated malignant metaprograms (**Figure 3a**) by performing Optimal Transport (OT) and assigning to each Visium HD cell the cell-type label of the most probable match from the snRNA-seq modality (see *Methods*). Visium HD-derived cell-type composition was consistent with snRNA-seq (Pearson correlation *r* = 0.47, 95% CI [0.24–0.64], *p-*value = 3.9×10^−6^), confirming overall agreement across platforms..

**Figure 3.**
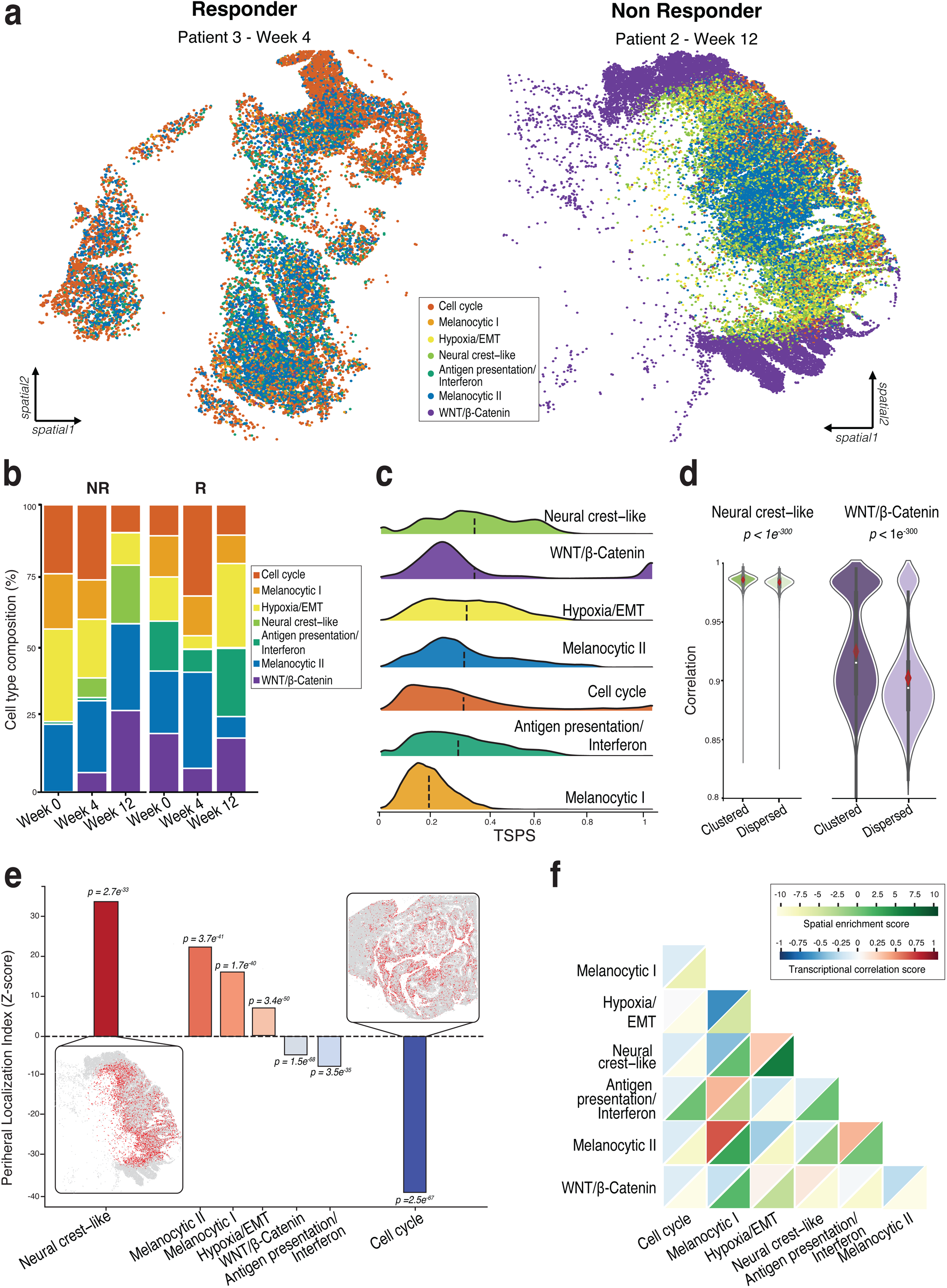
Analysis of the spatial organization of malignant cell states in Visium HD samples. **a.** Spatial arrangement of metaprograms (MPs) for a responder (left) and non-responder (right) patient. **b.** Bar plot of MP composition in NR and R patients across time points. **c.** Tumor State Proximity Score (TSPS) distribution of each MP. **d.** Pairwise correlations and statistical significance (two-sided Mann-Whitney U test p-value) between the expression profiles of Neural crest-like and Wnt/β-catenin cells stratified in *clustered* and *dispersed* groups based on their TSPS. **e.** Peripheral Localization Index (PLI) for each metaprogram, measuring the tendency of each cluster to localize near the tissue boundary (empirical permutation test, two-tailed p-value, 500 randomizations). Spatial maps of the MPs with the highest (*Neural crest-like*) and lowest (*Cell Cycle*) PLI are shown to illustrate boundary-enriched versus centrally enriched localization patterns. **f.** Spatial colocalization patterns (upper triangles) and transcriptional correlations (lower triangles) between MPs.

We first investigated tumor cell-intrinsic functional states in the context of their immediate malignant neighbors (homotypic interactions). Consistent with the RNA-seq modality, we observed a highly significant enrichment of *Neural crest-like* in NRs compared to Rs (Fisher’s exact test *p*-value < 0.05, odds ratio = 51.7, 95% CI [46.9, 57.1]) (**Figure 3b**). Conversely, *Antigen presentation/Interferon* was statistically enriched in Rs compared to NRs (Fisher’s exact test *p-*value < 0.05, odds ratio = 0.033, 95% CI [0.031, 0.035]). Spatial self-association of tumor states was captured using a Tumor State Proximity Score (TSPS, radius *r* = 500,see *Methods*). Briefly, the TSPS measures the spatial closeness between cells belonging to the same tumor state: higher values denoting spatial clustering and lower values reflecting cellular dispersion (**Figure 3c**). *Neural crest-like* formed the most compact clusters (TSPS mean = 0.35), consistent with its enrichment in NRs and suggesting the existence of a spatially consolidated treatment-resistant niche within such tumors. *Wnt/β-catenin* exhibited a rare bimodal TSPS distribution (TSPS mean = 0.34), indicating that in transcriptional programs either compact niches or dispersed cell subpopulations can spatially exist. *Antigen presentation/Interferon* showed a broad value distribution with moderate spatial clustering (TSPS mean = 0.28), whereas *Melanocytic I* displayed the most dispersed spatial arrangement (TSPS mean = 0.18). To test whether clustered homotypic interactions support coordinated cellular functions, following an approach similar to ^17^, we stratified Neural crest-like and Wnt/β-catenin cells according to their TSPS distributions (top ≥ 75th percentile as clustered, bottom ≤ 25th percentile as dispersed) and computed pairwise transcriptomic correlations within each subgroup. In both programs, clustered cells showed significantly higher pairwise correlations than dispersed cells, indicating that spatial clustering aligns with reduced transcriptional heterogeneity (**Figure 3d**).

To assess how spatial context may influence transitions between tumor phenotypes, we applied CellRank ^56^ along pseudotime to compute transition probabilities between spatial configurations of each cell state. Clustered cells were more likely to remain in the same state and maintain their spatial organization, while dispersed cells exhibited higher probabilities of transitioning to alternative states or spatial configurations (**Supplementary Figure 4**). Notably, Antigen presentation/Interferon and Hypoxia/EMT states showed the highest homotypic transition probabilities, highlighting their propensity for strong stabilization through spatial clustering.

Overall, these findings indicate that spatial clustering reinforces cell state identity, whereas dispersion enhances transcriptional plasticity and phenotypic instability.

### Spatial compartmentalization governs functional co-localization of tumor states

We then examined the composition of surrounding non-malignant cells (heterotypic interactions) to understand how the broader microenvironment shapes tumor cell states. We computed a Peripheral Localization Index (PLI) for each metaprogram, with negative values indicating peripheral enrichment, and positive values reflecting central localization (see *Methods*). Neural crest-like cells exhibited the highest central localization (PLI Z-score > 30, *p-*value = 2.7 × 10⁻³³, **Figure 3e**), alongside with the highest TSPS among all the MPs, indicating that these cells not only occupy the tumor core but also form densely clustered domains. This co-occurrence of central positioning and spatial compactness corroborated the notion that *Neural crest-like* represents a functionally cohesive, treatment-resistant subpopulation of cells. Other programs, including *Melanocytic I* and *II,* as well as *Hypoxia/EMT,* were enriched toward the tumor center, although to a lesser extent. In contrast, *Cell Cycle* was predominantly localized to the tumor periphery, with a markedly negative PLI Z-score (*p-*value = 2.5 × 10⁻⁶⁷), followed by *Antigen presentation/Interferon* and *Wnt/β-catenin*, which were similarly enriched at the tumor margins, albeit to a lesser extent.

We then examined the spatial colocalization and transcriptional correlations between malignant metaprograms as a measure of functional compartmentalization. We observed a positive association between *Melanocytic I* and *Melanocytic II*, indicating that these programs not only share similar melanocytic-like transcriptional identities but also occupy overlapping spatial niches (**Figure 3f**). Similar co-occurrence patterns were observed for *Hypoxia/EMT* with *Neural crest-like*, and *Melanocytic II* with *Antigen presentation/Interferon.* In contrast, *Cell Cycle* showed strong negative colocalization and transcriptional correlation with *Hypoxia/EMT* and *Wnt/β-catenin*, suggesting spatial segregation of these states within distinct tumor regions.

Collectively, these results indicate that malignant heterogeneity is organized not only by transcriptional divergence but also by spatial compartmentalization, with specific programs preferentially co-localizing to support divergent functional strategies within the tumor ecosystem.

### Spatial coordination of malignant and immune cells underlies response heterogeneity and ecosystem architecture

To comprehensively map the tumor ecosystem and its interactions with adjacent non-malignant cells (heterotypic interactions), we first annotated non-malignant subpopulations using the same OT methodology previously applied to malignant cells (**Figure 4a**). Consistent with the RNA-seq modality, the TME of the NR patients remained immunosuppressive throughout treatment, marked by persistent stromal elements and minimal immune infiltration (**Figure 4b**). In contrast, R patients exhibited a progressive increase in M1 macrophages, T cells, B cells, and plasma cells, reflecting a spatially active immune engagement and a shift toward antitumor immunity.

**Figure 4.**
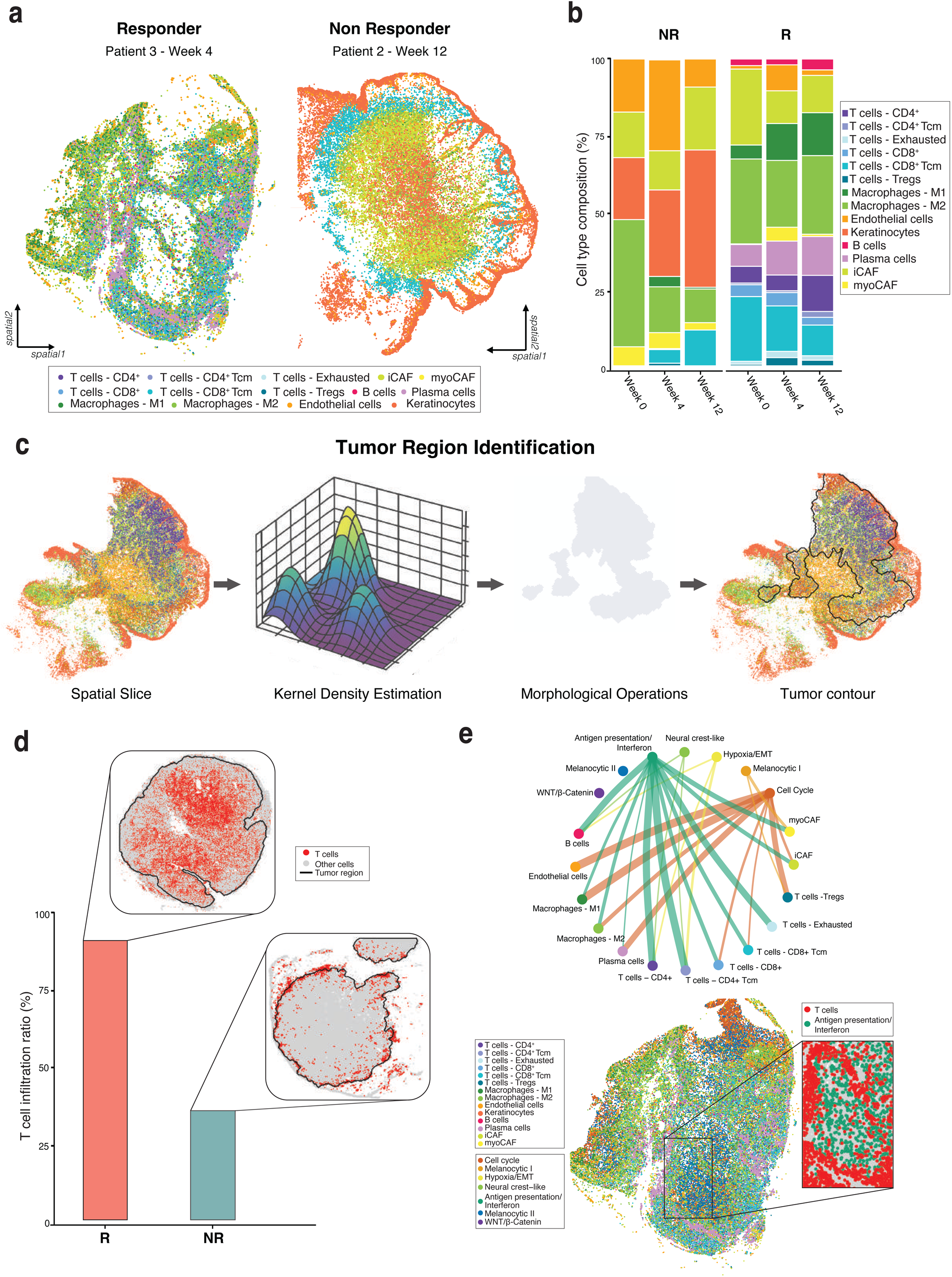
Analysis of the spatial organization of non-malignant cell states in Visium HD samples. **a.** Spatial arrangement of non-malignant cell states for a responder (left) and non-responder (right) patient. **b.** Bar plot of non-malignant composition in NR and R patients across time points. **c.** Computational pipeline for tumor region identification. Gaussian kernel density estimation (KDE) along with adaptive binarization threshold is followed by morphological operations (closing and dilation) to extract a valid tumor contour. **d.** T cell infiltration ratio (% of T cells inside the tumor region) between NR and R patients. **e.** Enrichment of malignant metaprograms (MPs) around non-malignant cell types. Top: MPs-non-malignant interaction for a R patient, where the width of the edge is proportional to the strength of the interaction. Bottom: spatial map highlighting colocalization of *Antigen presentation/Interferon* and T cells.

Then, tumor contours were delineated for each sample using a kernel density estimation (KDE) and morphological operations (**Figure 4c**; see *Methods*). Consistent with previous reports ^16,21^, we observed a significantly higher T cell infiltration in R compared to NR patients (**Figure 4d**). To investigate how malignant programs interact with their surrounding microenvironment, we quantified the local neighborhood composition of tumor cells, assessing which malignant states were preferentially enriched near distinct non-malignant cell types within a defined spatial radius. To distinguish true biological organization from random proximity, observed neighborhoods were compared against a permutation-based null model in which malignant program labels were shuffled while cell positions were preserved (see *Methods*). In R patients, the Antigen presentation/Interferon program strongly colocalized with B and T cells (**Figure 4e**, top), reflecting active immune response within these regions. This spatial proximity in a representative R patient (**Figure 4e**, bottom) highlights regions of close proximity between Antigen presentation/Interferon tumor cells and lymphocytes, indicative of enhanced antigen processing and presentation that promotes T and B cell activation and effective anti-tumor immunity.

To systematically capture recurrent spatial arrangements, we developed an unsupervised computational pipeline to define *ecotypes*, i.e. recurrent spatial composition patterns of cellular states across VisiumHD slides (see *Methods*). We identified five ecotypes named according to their cell type composition as *Antigen-presentation/Inflamed*, *Hypoxia*, *Immune–stroma interface*, *Keratinocyte epithelium*, and *Tumor core* (**Supplementary Figure 5a**). As shown in a representative specimen, Antigen-presentation/Inflamed ecotype dominated and intermixed with Immune–stroma interface regions, reflecting an activated immune microenvironment typical of R patients (**Supplementary Figure 5b)**. In contrast, NR tumors were dominated by Tumor core and Hypoxia ecotypes, coupled with stromal enrichment and limited immune infiltration (**Supplementary Figure 5c)**.

Together, these observations suggest that identical cellular components can assemble into divergent ecosystem architectures, distinguishing responsive, inflamed tumors from resistant, immune-excluded and stromal-dominated ones.

### Transposable Element reactivation drives immune signaling and contributes to tumor–immune dynamics

To evaluate the extent to which reactivated TEs contribute to shaping tumor and immune cell identities, as well as their interactions, we quantified TE activity across both malignant and non-malignant compartments. Leveraging a multimodal integrative framework optimized for locus-specific quantification of TEs in single-cell sequencing ^57^, we profiled the expression and chromatin accessibility of major TE classes, including LINEs, SINEs, LTRs, and other repetitive elements. TE-derived signals proved highly informative for resolving the cellular architecture of the TME, with co-embedding t-SNE analysis separating stromal, T cell, and macrophage sub-populations (**Figure 5a**, left). Moreover, TE activity reflected malignant cell heterogeneity across transcriptional MPs, revealing a continuum of tumor cell states rather than sharply segregated clusters (**Figure 5a**, right).

**Figure 5.**
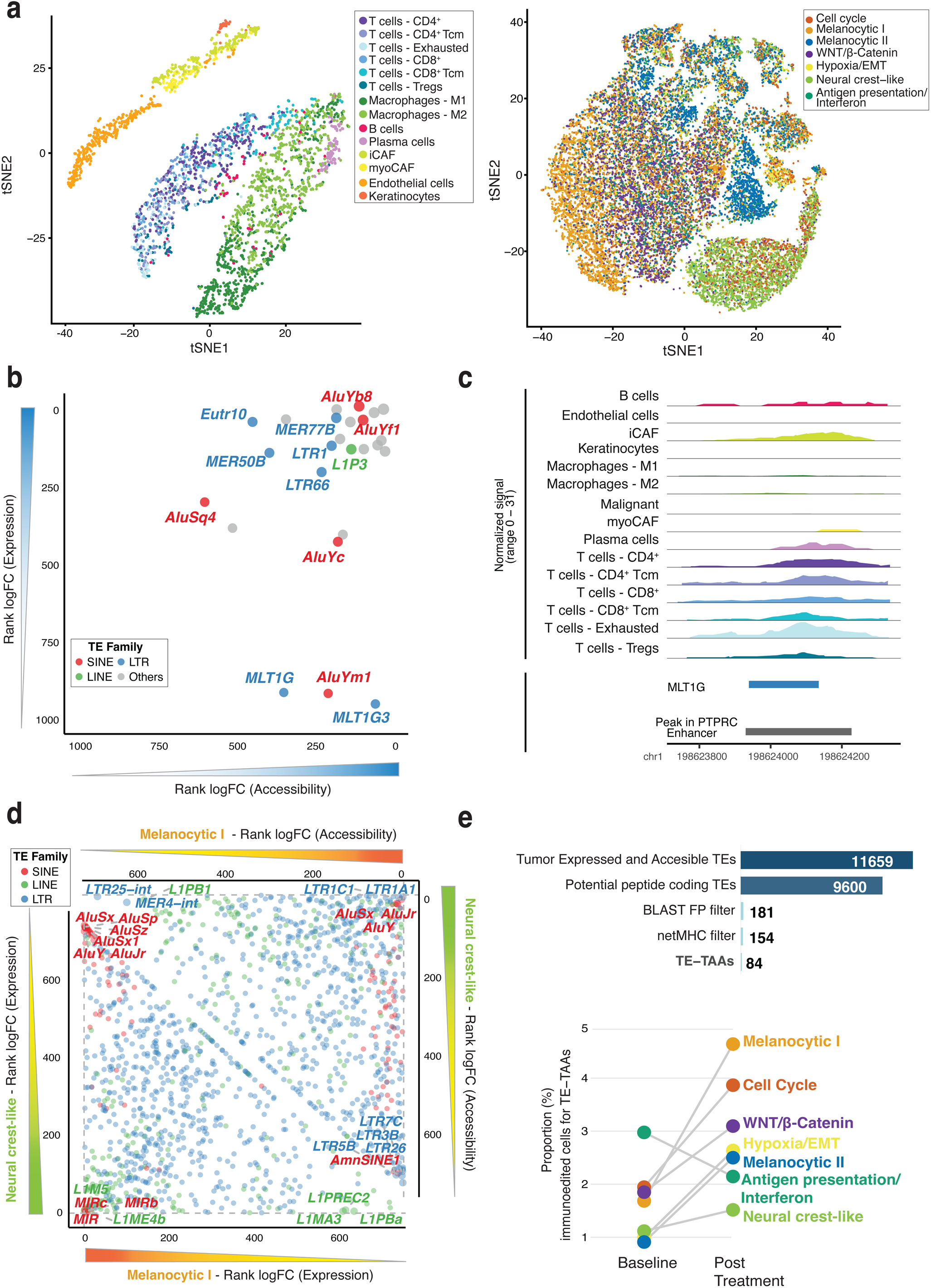
TE activity across tumor and TME states under epi-immunotherapy. **a**. t-SNE of co-embedding of transposable elements (TEs) from paired snRNA-seq and snATAC-seq across TME cells (left) and tumor metaprograms (right). **b.** Differential analysis of TE activity in TME cells pre-therapy versus on-treatment (week 4), comparing chromatin accessibility (x-axis) and expression (y-axis), ranked by log_2_FC; points colored by TE family. **c.** Accessibility of an MLT1G locus overlapping a *PTPRC* enhancer, showing normalized accessibility signal across all TME subtypes and malignant cells. **d**. Comparative analysis of TE activity between *Melanocytic I* and *Neural crest-like* states, highlighting specific and shared TE patterns. Upper triangle: chromatin accessibility fold-change ranks; lower triangle: expression fold-change ranks; features colored by TE family. **e.** Top: filtering strategy and number of TE-derived tumor-associated antigens (TE-TAAs). Bottom: fraction of malignant cells classified as immunoedited, defined by concurrent TE expression and accessibility at loci generating patient-matched immunogenic TE-TAAs. Percentages of immunoedited cells were calculated per metaprogram and compared across treatment time points.

We next assessed whether therapeutic intervention with demethylating agents modulated TE dynamics in the TME. Differential analysis of TME cells before therapy (week 0) versus on-treatment (week 4) revealed that DNMTi increased TE activity, evident at both the level of chromatin accessibility (x-axis) and transcriptional expression (y-axis) for specific TE elements (**Figure 5b** and **Supplementary Table 9**). A subset of TEs showed simultaneous high accessibility and expression after treatment, including SINE Alu elements (e.g., *AluYb8* and *AluYf1*), LTR elements (e.g., *MER77B* and *LTR1*), and LINE-1 elements (e.g., *L1P3*), which emerged as top-ranking features in both RNA and accessibility analyses. Notably, Alu elements represent some of the most evolutionarily recent and transcriptionally active TEs in the human genome (mean ages of 1.89 and 5.36 Myr, respectively), suggesting that DNMTi treatment preferentially reactivates young, transcriptionally competent elements. In contrast, different TE families, such as LTR elements as *MLT1G* and *MLT1G3*, exhibited reactivation predominantly at the accessibility level, without corresponding transcriptional upregulation. This pattern suggested a potential *cis*-regulatory role of such elements, whereby epigenetic reprogramming induced by DNMTi may prime these elements to act as regulatory enhancers, contributing to the transition of immune cells from a resting to an activated state in response to treatment. We indeed observed that a specific *MLT1G* locus (chr1: 198,623,845–198,624,639, GRCh38/hg38) overlapped an accessible chromatin region annotated as a PTPRC (CD45) enhancer and an active regulatory feature in CD45^+^ cells (**Figure 5c**).

To dissect the contribution of TEs to malignant identities and transcriptional programs, we identified differentially expressed and accessible TEs across MPs (**Figure 5d** and **Supplementary Table 10**). TE involvement varied along the melanoma differentiation axis, from the highly differentiated Melanocytic I to the stem-like, de-differentiated Neural crest-like state. While some TEs, including *LTR1C1*, *LTR1A1*, and *AluSx*, *AluJr*, *AluY*, were accessible across both states, the Alu elements, known for their role in viral mimicry and activation of innate immune sensing that may enhance responsiveness to epigenetic therapy ^58^, were selectively re-expressed in *Melanocytic I* tumor cells.

We then assessed whether re-activated TEs could also serve as a source of tumor-associated antigens (TE-TAAs), which are known to enhance immune responses under epigenetic interventions ^59^. Through a stringent computational workflow, we identified 84 high-confidence TE loci with potential to generate TAAs, corresponding to 154 unique TE-derived sequences with predicted antigenicity (**Figure 5e, top** and **Supplementary Table 11**). To quantify the fraction of malignant cells potentially subject to immune elimination through TE-TAAs, we defined as *immunoedited* those cells that simultaneously (i) displayed expression and chromatin accessibility at a TE locus potentially generating a TE-TAA, and (ii) matched an immunogenic TE-TAA identified in the corresponding patient. For each MP, we then calculated the percentage of immunoedited cells relative to the total malignant cells of that MP and compared their abundance across treatment time points (baseline at week 0 versus post-treatment at weeks 4 and 12). Strikingly, only tumor cells belonging to the Antigen presentation/Interferon state, previously characterized by strong antigen-presentation machinery and immunoproteasome gene expression, declined following treatment, consistent with a potential immune-mediated clearance driven by TE-derived antigens (**Figure 5e, bottom**). Notably, two TE-TAA–derived epitopes, *RPPLRPANFLY* (from the Alu family) and *KYGPNSPYM* (from the human ERVK family), have been independently reported as highly immunogenic ^60,61^.

Altogether, these results highlight a dual role for TEs in melanoma under epi-immunotherapy: acting in *cis* as potential regulatory elements shaping immune activation, and in *trans* by generating immunogenic peptides that contribute to tumor–immune interactions, modulating both cell identity and tumor plasticity. Moreover, these results support the notion that one of the main mechanistic effects of the epigenetic therapy is to promote the transcription of TE-derived antigens, and enhance anti-tumor immune response as previously observed in glioma ^62^ and lung ^63^.

### NFATC2 drives resistance to ICI and epi-immunotherapy, and its targeting shifts Neural crest–like melanoma towards a differentiated phenotype

Melanoma cells can shift from proliferative/differentiated states to invasive/de-differentiated states, the latter being linked to poorer response to ICI therapy and reduced survival ^64^ ^40,65^.

To assess the clinical relevance of the transcriptional MPs identified in our single-cell analyses, we analyzed a large collection of eight previously published melanoma patient cohorts. These patients were treated with different ICI treatments, and their tumors were profiled by bulk transcriptomics before and/or after therapy. To minimize confounding effects from non-malignant cells, we retained only high-purity tumor samples (purity ≥ 0.3, n = 402, R = 152, NR = 250).

Using a deconvolution approach based on MP-specific gene signatures ^66^, we estimated the proportion of malignant cells corresponding to each MP in the bulk tumor samples, and evaluated differences according to the response group (**Figure 6a**). Consistent with our findings and prior observations ^16^, tumors from Rs were significantly enriched for the Antigen presentation/Interferon state both before and after ICI treatment (Wilcoxon test, *p-*value = 0.003 and 0.031, respectively), highlighting its predictive value for therapy response. Conversely, we found that the Melanocytic I/II states were more abundant in NRs prior to treatment (Wilcoxon test, *p-*value = 0.017 and 0.025, respectively). Notably, the de-differentiated *Neural crest-like* cells became specifically enriched in NRs after treatment (Wilcoxon test, *p-*value = 0.015). Intriguingly, the relative abundance of these two distinct cell populations at single-cell resolution, the Antigen presentation/Interferon and Neural crest-like states, demonstrated strong diagnostic performance in predicting therapy response (**Supplementary Figure 6a**). Their predictive power was notably enhanced in the context of epi-immunotherapy compared to ICI alone, indicating that different therapeutic combinations may target or be more effective against distinct malignant cell states. Importantly, a higher fraction of Neural crest-like cells was significantly associated with poorer overall survival, both pre- and post-ICI treatment (Hazard ratio = 0.67 and 0.50, respectively; *p-*value < 0.05), reinforcing its prognostic significance (**Figure 6b** and **Supplementary Figure 6b**).

**Figure 6.**
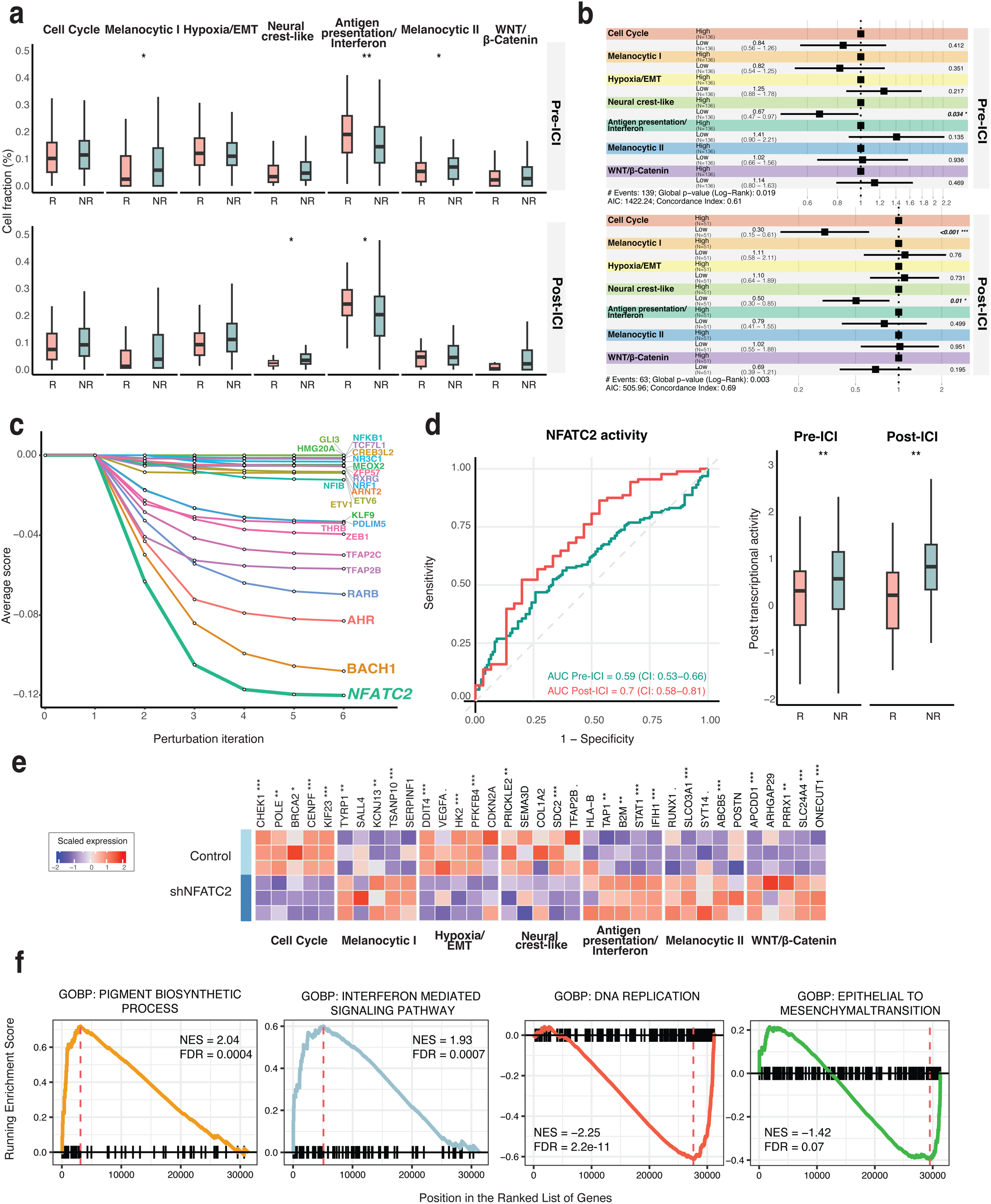
Prognostic value of metaprograms and regulatory role of NFATC2. **a.** Boxplot showing MP deconvolution in bulk RNA-seq cohorts stratified by pre- and post-ICI treatment (pre-ICI: R = 30, NR = 88; post-ICI: R = 122, NR = 162). Statistical significance was assessed using Wilcoxon rank-sum test; * *p* ≤ 0.05, ** *p* ≤ 0.01. **b.** Forest plot showing overall survival (OS) by MP deconvolution in bulk RNA-seq cohorts, pre- and post-ICI treatment. Patients were grouped as High or Low based on median MP abundance. HR (95% CI) calculated using Cox regression. **c.** Predicted changes in the Neural crest-like transcriptional state following simulated knockdown of MP-associated TFs. For each TF, the mean difference in Neural crest-like signature gene expression across five independent iterations is shown. **d.** Receiver operating characteristic (ROC) curves of response prediction in pre- and post-ICI based on NFATC2 post-transcriptional activity (left), and boxplots of activity scores stratified by responder status (R vs NR) before and after ICI (right). Areas under the ROC curve (AUCs) are shown with 95% confidence intervals (CIs). Statistical significance was assessed using the Wilcoxon rank-sum test; ** p ≤ 0.01. **e.** Heatmap of scaled expression of selected genes in control and shRNA NFATC2 conditions. Rows represent conditions and columns represent marker genes grouped by MP. Statistical significance of differential expression was assessed with limma; . p ≤ 0.1, * p ≤ 0.05, ** *p* ≤ 0.01, *** *p* ≤ 0.001. **f.** Enrichment plots of representative GOBP terms after NFATC2 knockdown.

Our previous finding showed that melanoma *Neural crest-like* cells exhibit a lower level of plasticity than other melanoma cells; therefore, we explored potential molecular drivers of these therapy-resistant transitions. We performed a computational perturbation analysis using SCENIC+ ^67^ to determine how perturbing the expression of key master regulator transcription factors would impact this cell state (**Figure 6c**). Among the candidates, simulated silencing of *NFATC2* (Nuclear Factor of Activated T-cells, cytoplasmic 2) produced the most pronounced effect, nominating it as a central regulator of the resistant Neural crest-like state.

To characterize *NFATC2* function, we inferred its regulon from the independent single-cell dataset of melanoma patients used before ^16^. This approach was crucial for minimizing potential bias from our own epigenetic therapy dataset, allowing us to define a high-confidence NFATC2 regulon, consisting of 56 genes (**Supplementary Figure 6c**). As expected for a program sustaining a Neural crest-like phenotype, this regulon contained a number of genes associated with embryonic development, invasion, migration, and neurogenesis. Among these genes, we found key drivers of developmental mechanics, such as the AP-2 family members *TFAP2C* and *TFAP2B*, which are critical for the development of ectodermal tissues and the neural crest ^68^. We also identified genes that are key drivers of tumor cell invasion and migration, including *CSPG4* and *CAV1*, genes previously associated with tumor cell migration, poor prognosis and multidrug resistance in cancer ^69,70^. As anticipated, the regulon also contained genes with established roles in neurogenesis and neuronal function, such as *MAP2* and *SEM3D,* markers of neuronal differentiation and cell migration and axon growth ^71,72^. The transcriptional activity of the *NFATC2* regulon resulted in a powerful predictor of secondary resistance to ICI therapy (AUC = 0.7, **Figure 6d**), with significantly higher activity in NR samples both before and after therapy (Wilcoxon test, *p*-value = 0.007 and 0.001, respectively).

While previous studies have already linked NFATC2’s target gene signatures to ICI resistance in melanoma ^73^, our findings directly associate *NFATC2* in driving the Neural crest-like phenotype itself, providing a mechanistic link between cell state and therapeutic failure. These results are consistent with prior studies showing that, in MITF^low^ melanomas, *NFATC2* regulates the EMT-like transcriptional program, invasive potential, and tumor growth in vitro and in vivo, in part via oncogenic networks involving *c-MYC, FOXM1,* and *EZH2* ^74–76^.

To connect our in silico findings with experimental evidence, we analyzed stable *NFATC2* knockdown data in a melanoma cell line derived from a surgical specimen ^74–76^. By overlapping the top 50 DEGs of each MP (**Supplementary Table 4**) with DEGs from *NFATC2*-silenced cells (**Supplementary Table 13**), we found robust inactivation of both Cell Cycle and Neural crest-like programs (**Supplementary Figure 6d**). Key downregulated genes included *CHEK1*, *POLE*, and *CENPF* (*Cell Cycle*) and *PRICKLE2*, *SDC2*, and *TFAP2B* (*Neural crest-like*) (**Figure 6e**). Gene set enrichment confirmed suppression of DNA replication and EMT transitions, accompanied by an increase of interferon mediated signalling and pathways associated with melanocyte differentiation, such as pigment biosynthetic process (**Figure 6f** and **Supplementary Table 14**). Conversely, we observed a transcriptional shift toward a more differentiated *Melanocytic II* state (Fisher’s method, *p*-value = 4.4 × 10^−4^, **Supplementary Figure 6d**), with reactivation of pigmentation pathways, and simultaneously activated interferon-mediated signaling, including upregulation of specific genes of the *Antigen presentation/Interferon* MP such as *TAP1*, *B2M*, *STAT1*, and *IFIH1*.

Together, these findings establish *NFATC2* as a key regulator of the Neural crest–like melanoma state, coupling proliferative and migratory programs with therapy resistance. Silencing of *NFACT2* not only reverted this resistant state but also promoted differentiation and immunogenic reprogramming, highlighting *NFATC2* as a potential therapeutic vulnerability in aggressive melanoma. Importantly, the *NFATC2*-driven program may also contribute to mechanisms of resistance against epi-immunotherapy, positioning its inhibition as a strategy to enhance responsiveness to such treatments.

## Discussion

Our multi-omic and spatial dissection of longitudinal tumor biopsies provides a mechanistic framework for how epi-immunotherapy reshapes the melanoma ecosystem. We identify tumor cellular states and regulatory programs that underlie therapeutic sensitivity and resistance, revealing actionable vulnerabilities. Prior bulk and single-cell studies established the existence of discrete melanoma cellular states, such as MITF^high^ melanocytic, AXL^high^ invasive, stress-related, antigen-presenting, and neural crest-like, linked to therapy response and resistance in ICI therapies ^1,4,14,15^. We confirm these states but, crucially, show how they evolve dynamically under combined DNMT1 inhibition and CTLA-4 blockade. In particular, we uncover substantial heterogeneity within the melanocytic compartment, resolving three distinct transcriptional programs. One of these programs is uniquely characterized by activation of the Wnt/β-catenin signaling pathway, evidenced by strong expression of *LEF1*. This represents a previously unappreciated melanocytic sub-state with potential relevance for immune evasion and therapeutic resistance ^77^. The persistence of a Neural crest-like program in non-responders and enrichment of an Antigen presentation/Interferon program in responders parallels recent observations of phenotype switching as a mechanism of immune evasion ^16^, while adding mechanistic depth from paired transcriptome–chromatin profiling.

Our study highlights the critical role of spatial organization in shaping melanoma cellular states and their functional stability under epi-immunotherapy. By integrating TSPS analysis with trajectory-based transition modeling, we demonstrate that clustered tumor cells exhibit both greater transcriptional homogeneity and higher probability of maintaining their state and spatial configuration compared to dispersed counterparts. These findings indicate that spatial clustering reinforces cell state identity, whereas dispersion promotes transcriptional heterogeneity and state instability. This principle has been recently described in other cancer contexts ^17,78^, while studies of circulating tumor cell clusters have demonstrated that homotypic adhesion supports survival and functional stability during metastasis ^79,80^. Collectively, these observations support a concept in which differential homotypic adhesion acts as an organizing principle in cancer, preserving cellular identity in clustered niches and enabling plasticity in dispersed cells. The unusual bimodal TSPS distribution of the Wnt/β-catenin program provides a further example of the relationship between pathway biology and spatial architecture. β-catenin, at adherens junctions, stabilizes cadherin-mediated homotypic adhesion, fostering cohesive architecture; whereas in the nucleus, it functions as a transcriptional co-activator, driving programs linked to plasticity, invasion, and immune evasion ^81^. The coexistence of clustered and dispersed Wnt/β-catenin subpopulations may therefore reflect these two roles of β-catenin. Clustered cells likely exploit adhesion-mediated stabilization to maintain coherence, while dispersed cells may engage β-catenin’s transcriptional activity, resulting in greater heterogeneity and state transitions. Interestingly, we also observed that Antigen presentation/Interferon and Hypoxia/EMT programs displayed the highest homotypic transition probabilities. These states may represent functionally reinforced niches, where spatial clustering enhances stability and ensures persistence under therapeutic pressure. Conversely, dispersed cells across states showed lower self-transition probabilities, consistent with their role as reservoirs of plasticity and adaptation. Taken together, our findings reveal a model in which spatial clustering and dispersion constitute parallel organizational strategies in melanoma: clustered niches, stabilized by homotypic adhesion, preserve state identity and may contribute to treatment resistance, while dispersed cells provide a source of heterogeneity and adaptive potential.

Our results support and extend prior work showing that epigenetic drugs can enhance responsiveness to immune checkpoint inhibitors by inducing viral mimicry and innate immune signaling ^12,82–84^. We demonstrate that TE reactivation has a dual role: acting as *cis*-regulatory elements that prime immune activation, and in *trans* generating TE-derived antigens cleared by T cells. While similar TE-mediated immunogenicity has been reported in other cancers ^62,63^, our study provides single-cell resolution evidence that TE-driven antigenicity actively shapes melanoma cell fate under epi-immunotherapy. Importantly, we uncover an inverse association between TE reactivation and the *Antigen Presentation/Interferon state*, supporting the hypothesis that tumor immuno-editing in the context of epigenetic therapy is highly dependent on such cellular state. This highlights TE reactivation not merely as a global effect of treatment, but as a state-specific mechanism that contributes to immune pressure and fate decisions in melanoma. Prior studies have implicated NFATC2 in EMT-like transitions and resistance in MITF^low^ melanoma ^73,75,76^. We identify NFATC2 as a master regulator of a specific therapy resistant cellular state linking invasive behavior and immune evasion. This positions NFATC2 inhibition as a rational therapeutic strategy to overcome both primary and acquired resistance to epi-immunotherapy.

One of the main limitations of our study consists of the small number of patients and limited sampling time points/technical challenges, like obtaining sufficient high-quality tissue inherent to/proof-of-concept trials; therefore, larger studies will be required to generalize some of our observations ^85^. Moreover, functional validation of candidate regulators such as NFATC2 in vivo will be essential to establish causality. Nevertheless, together, our results establish a framework in which epi-immunotherapy reshapes melanoma ecosystems by promoting immune-engaged states while exposing vulnerabilities of resistant niches. Through integration of single-cell multiome and spatial analyses, we provide mechanistic insight into tumor plasticity, TE-mediated immune dynamics, and regulatory drivers of resistance.

## Methods

### Experimental method details

#### Sample collection

Snap-frozen tumor tissues (*n* = 13) were collected from five patients with advanced cutaneous melanoma enrolled in the phase Ib NIBIT-M4 clinical trial (NCT02608437), as previously described ^11^. Briefly, patients received escalating doses of the DNA hypomethylating agent guadecitabine (30–60 mg/m²/day, subcutaneously on days 1–5) in combination with the standard anti-CTLA-4 ipilimumab (3 mg/kg, intravenously on day 1) across four treatment cycles. The study was conducted in accordance with the Declaration of Helsinki and Good Clinical Practice guidelines, with protocol approval by the ethics committee of the University Hospital of Siena. All patients provided written informed consent. Tumor biopsies for correlative analyses were collected longitudinally at baseline (week 0), during treatment (week 4), and at a later on-treatment time point (week 12). Patients were classified as Responder (R; complete response [CR], partial response [PR], or stable disease [SD]) or Non-Responder (NR; progressive disease [PD]) patients, based on clinical response.

#### 10X Genomics Single-Nucleus Multiome (RNA + ATAC) sequencing

Nuclei were isolated from 13 snap-frozen tissues using the Nuclei isolation kit (10x Genomics, CG000505 Rev A) following the manufacturer’s instructions. Nuclei counts and viability were assessed using Nexcelom Cellometer K2 (Nexcelom Bioscience). Within a range of 3,000 - 10,000 nuclei per sample were used for Single cells multiome libraries generation according to the Chromium Next GEM Single cell Multiome ATAC + Gene Expression (10x Genomics, CG000338 Rev F). Briefly, transposed nuclei were combined with barcoded Gel Beads, a Master Mix, and Partitioning Oil on a Chromium Next GEM Chip J for gel bead-in-emulsions (GEM) generation and barcoding. GEX and ATAC libraries were processed separately and their quality was examined on 5200 Fragment Analyzer (Agilent Technologies). Sequencing was performed on Novaseq X Plus (Illumina).

#### Visium HD spatial gene expression profiling

Formalin-fixed paraffin-embedded (FFPE) tissues were reviewed by a board-certified pathologist and placed on the Visium slide following the Visium CytAssist Spatial Gene Expression for FFPE – Tissue Preparation Guide (10x Genomics, CG000518, Rev C). Spatial gene expression libraries were generated following the Visium HD Spatial Gene Expression Reagent Kits (10x Genomics, CG00685, Rev A). Sequencing was performed on Novaseq X Plus (Illumina).

### Data processing and statistical analyses

#### Single-Nucleus Multiome data processing

The 10x Genomics Cell Ranger ARC (v2.0.2) pipeline was used to process the multiome data. The GRCh38 human reference was used for the read mapping (refdata-cellranger-arc-GRCh38-2020-A-2.0.0). The raw RNA count matrix and ATAC fragment data were further processed using R packages Seurat (v5.1.0) ^86^ and Signac (v1.14.0), respectively ^87^.

#### RNA-seq modality

Low-quality nuclei were excluded from the RNA-seq dataset based on standard quality control (QC) metrics. Specifically, for each sample, nuclei exhibiting either low or excessively high gene complexity (< 500 or > 7,500 detected genes), elevated mitochondrial gene content (> 5%, indicative of dying or stressed cells), or abnormally high total transcript counts (nCount > 50,000) were filtered out. Genes detected in at least three nuclei were retained. After QC filtering, a total of 58,670 high-quality nuclei were retained for downstream analyses (median 3,675 nuclei per sample, with 5,722 unique molecular identifiers [UMIs] and 2,774 genes detected per nucleus).

The merged gene expression matrix was subsequently normalized and log-transformed. Dimensionality reduction was performed via principal component analysis (PCA) using the top 3,000 highly variable genes. The first 22 principal components were then used to construct a t-distributed stochastic neighbor embedding (t-SNE) for visualization, and unsupervised clustering was carried out using a resolution of 0.2.

#### ATAC-seq modality

Raw ATAC-seq fragment data were processed by merging the common peaks across samples using the *reduce* function from the GenomicRanges package (v1.54.1). Peaks shorter than 20 or longer than 10,000 base pairs were excluded as outliers.

Nuclei with low-quality chromatin accessibility profiles were excluded based on standard QC criteria. Specifically, nuclei with fewer than 2,500 or more than 20,000 total fragments were removed. Additional exclusion criteria included a low fraction of fragments in peaks (percent reads in peaks < 20%), a high proportion of reads mapping to ENCODE blacklisted regions ^88^ (blacklist ratio > 0.05), elevated nucleosome signal (> 1.5), and low transcription start site (TSS) enrichment (< 2). Following QC filtering, a total of 55,850 high-quality nuclei were retained for downstream analysis (median of 3,944 nuclei per sample, with 16,472 fragments per cell and a TSS enrichment of 3.83).

The resulting peak count matrix was normalized using term frequency–inverse document frequency (TF-IDF) transformation. Dimensionality reduction was performed via singular value decomposition (SVD) on the top 95% most commonly accessible peaks (*FindTopFeatures with min.cutoff set to ‘q5’*). The latent semantic indexing (LSI, 2:50) components from the SVD reduction were used for secondary dimensionality reduction via t-SNE. The first LSI component, which was strongly correlated with sequencing depth, was excluded from downstream analyses. Unsupervised clustering was performed using the smart local moving (SLM) algorithm implemented in the Seurat package (*FindClusters* with *algorithm* = 3, at a resolution of 0.8). Peak calling was conducted using the MACS2 package (v2.2.6) ^89^ via the *CallPeaks* function. Gene activity scores were computed using the *GeneActivity* function, which aggregates chromatin accessibility across gene bodies and promoter regions.

#### Integrating modalities

To integrate information from both RNA and ATAC modalities for multiomic clustering, nuclei passing quality control filters in both datasets (n = 38,029) were retained. The *FindMultiModalNeighbors* function in Seurat was used to construct a Weighted Nearest Neighbor (WNN) graph by jointly considering the RNA Principal Components (PCs 1:14) and ATAC latent semantic indexing components (LSI 2:50) ^90^. This WNN graph was used as input for dimensionality reduction using t-SNE. Unsupervised clustering was subsequently performed on the Weighted Shared Nearest Neighbor (WSNN) graph using the SLM algorithm with a resolution parameter set to 0.1.

#### Malignant and non-malignant cell discrimination and tumor microenvironment (TME) cell type annotation

Classification of cells into malignant and non-malignant cells was performed individually on RNA modality using SCEVAN package (v.1.0.3) ^18^, which integrates single-cell expression data with inferred copy number alterations to identify malignant clones. Following malignancy assignment, non-malignant cells were clustered using the first 12 PCA and a resolution of 0.7. Subsequently, cell types were annotated using well-established immune and stromal cell-type marker signatures ^91^, combined with the inspection of canonical marker genes at both the gene expression and chromatin accessibility levels. To systematically evaluate immune cell-type enrichment across clusters, we applied the Normalized Enrichment Score (NES) derived from the Mann-Whitney-Wilcoxon Gene Set Test (mww-GST), as previously described in ^92^ on gene ranks obtained by differential expression analysis between each Seurat cluster and all others. Based on this integrative analysis, the TME was resolved into distinct cellular subtypes, including components of the immune compartment: T cells (CD8^+^, CD4^+^, exhausted, and regulatory T cells), B cells, plasma cells, and macrophages (M1 and M2 subtypes), as well as the stromal compartment, comprising cancer-associated fibroblasts (inflammatory CAFs [iCAFs] and myofibroblastic CAFs [myoCAFs]), endothelial cells, and keratinocytes.

#### Malignant cell meta-programs identification

Malignant cell meta-programs (MPs) were identified following the approach described by ^21^. Briefly, non-negative matrix factorization (NMF) was independently applied to the relative gene expression profiles of malignant cells from samples containing at least 200 malignant cells. Prior to NMF, all negative expression values were set to zero. To avoid predefining the number of transcriptional programs (rank k), NMF was run across multiple values (k = 4, 5, …, 9), yielding a total of 468 NMF programs. Each NMF program was defined by the top 50 genes with the highest NMF coefficients. To ensure biological relevance and consistency, only robust and non-redundant NMF programs were retained. Robustness was defined by high intra-sample consistency (minimum 70% gene overlap across different k values), low redundancy within samples (maximum 20% overlap), and limited overlap across patients (maximum 20%). This filtering resulted in 75 robust NMF programs. These programs were subsequently grouped based on pairwise Jaccard similarity using an iterative hierarchical clustering approach that merged highly overlapping programs, as previously described ^21^. Each resulting MP was defined by the 50 most recurrent genes across the NMF programs within each cluster.

#### Classification of malignant cells to MP states and functional characterization

Each malignant cell was scored for all seven MP states using a gene set scoring approach. Specifically, the *AddModuleScore* function from the Seurat package was used to compute a signature score for each MP based on its corresponding 50 gene set. Cells were retained for further classification only if their highest MP signature score exceeded 0.3.

To quantify the confidence in assigning a cell to a specific MP state, a *simplicity score* was computed as follows:

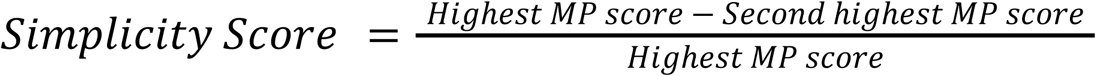

Cells with a simplicity score ≥ 0.1 were confidently assigned to the MP corresponding to their highest score. To assess the biological relevance of each MP, functional enrichment analyses were applied. Overrepresentation analysis (hypergeometric test) was conducted on the 50-gene MP signatures. In parallel, Gene Set Enrichment Analysis (GSEA) was performed using the clusterProfiler package (v4.10.1) ^93^ on ranked gene lists derived from differential expression between each MP and all other malignant cells, as computed with the memento framework (v0.1.1) ^94^. Curated gene sets were obtained from Gene Ontology (GO), Reactome, KEGG, Hallmark, and WikiPathways databases. Additionally, differentially expressed genes (DEGs) for each MP were defined using the thresholds of fold change (de_coef ≥ |ln(2)|) and adjusted *p*-value < 0.05.

Cell-cell communication analysis was performed to evaluate interactions between malignant cells (stratified by MP assignment) and tumor microenvironment (TME) cell populations using the CellChat package (v2.1.2) ^95^.

#### Cross-modal concordance analysis

Concordance between transcriptional and chromatin accessibility states was assessed by performing cross-modal label transfer with Seurat (v5.1.0) ^86^. The snRNA-seq dataset was used as the reference and gene activity scores from the snATAC-seq dataset were used as the query. Cross-modal anchors were identified using canonical correlation analysis (CCA) with *FindTransferAnchors*, and labels were transferred with *TransferData* (LSI 2:50). Predicted annotations were assigned to individual snATAC-seq nuclei, compared with their reference states, and prediction scores were averaged across cell types to evaluate correspondence between transcriptional identities and accessibility-based predictions.

#### Melanoma related Signature

Transcriptional metaprograms were compared with published melanoma gene signatures ^4,14,16^. Module scores in malignant cells were quantified using the Seurat function *AddModuleScore*, and concordance with published signatures was assessed by Pearson correlation analysis.

#### Transcription factor motif enrichment analysis

Transcription factor (TF) motif enrichment was analyzed on differentially accessible (DA) peaks for each metaprogram relative to all others (average log_2_FC ≥ 0.58 and adjusted *p-* value < 0.05). To mitigate sequence composition bias, accessible peaks were matched to background regions with comparable GC content using *MatchRegionStats* function from Signac. Enrichment was tested with *FindMotifs* hypergeometric framework from Signac, using TF position weight matrices from the HOCOMOCO v13 database. Motifs with fold enrichment ≥ 1.3 were retained. To avoid redundancy, TFs represented by multiple motifs were filtered such that only the most strongly enriched motif per TF was kept. The union of top 30 TFs per metaprogram, along with *MIFT*, was displayed as a heatmap, with values representing fold enrichment over background. Motif footprinting for selected TFs was performed with the *Footprint* function from Signac using the GRCh38/hg38 genome assembly.

#### Cis-regulatory region (CRE)-TFs-gene linkage analysis

Genes linked to enriched TF motifs (as described above) were mapped to promoter-like signatures (PLS), proximal enhancer-like signatures (pELS), and distal enhancer-like signatures (dELS) based on ENCODE candidate cis-regulatory element (cCRE) annotations ^96^. For each regulatory category, motif-containing cCREs were intersected with their linked protein-coding genes, and only those also identified as up-regulated from DEG analysis between each MP vs. the others were retained. These filtered gene sets were then subjected to over-representation analysis (ORA) using the *enricher* function from clusterProfiler ^93^, using curated gene sets from databases including Gene Ontology (GO) biological process categories, Reactome, KEGG, and Hallmark.

#### Enhancer-driven gene regulatory network analysis

Enhancer-driven gene regulatory networks (eGRNs) were inferred from snRNA-seq and snATAC-seq multi-ome data using SCENIC+ (v1.0a2)^67^. First, pycisTopic (v2.0a0) was used to identify and binarize cis-regulatory topics from the snATAC-seq dataset, and pycisTarget (v1.1) was used to identify enriched motifs. The SCENIC+ Snakemake pipeline was executed with default parameters to predict enhancer-driven regulons constituting the eGRN. Regulon specificity scores (RSS) were calculated for each MP. High-quality eRegulons were defined by computing the correlation between TF expression and the activity scores (AUC) of their predicted target regions, retaining only TF–region pairs with correlation coefficients above |0.3| yielded 188 eRegulons with a median of 196 genes and 292 regions each. eRegulon activity was quantified using AUCell scores ^97^. TFs were subsequently ranked based on: (i) TF expression in a given MP relative to all other MPs, (ii) eRegulon AUCell scores on gene expression in a given MP relative to all other MPs, and (iii) eRegulon AUCell scores on accessibility in a given MP relative to all other MPs. Only TFs ranked within the top 30 across all three criteria (n unique TFs = 104) were retained for downstream analyses (**Supplementary Table 7**).

#### Visium HD data processing

##### Nuclei Segmentation and Binning

We performed nuclei segmentation on the high-resolution hematoxylin and eosin (H&E) stained images to delineate individual nuclei, enabling the assignment of spatial barcodes to specific cells. In detail, a pre-trained deep learning model, StarDist ^98^, was used to create a nuclei mask; the shape and location of each nucleus in the mask were then employed to sort barcodes into bins corresponding to each individual nucleus. Finally, we aggregated gene expression counts from the 8×8 *µ*m barcoded squares associated with each segmented nucleus by performing gene-wise summation of UMI counts, resulting in a gene-by-cell matrix approximating single-cell resolution. This procedure was independently repeated for each of the 9 samples, resulting in an average of 113,366 cells per sample with 18,085 genes.

##### Data preprocessing

For each sample, low quality cells and genes were removed. Specifically, we filtered out cells showing high mitochondrial gene content (>5%) or expressing fewer than 5 detected genes. Additionally, genes expressed in fewer than 5 cells were also removed to reduce noise and focus on biologically relevant signals. A total of 736,028 cells passed the quality control step and their gene expression counts were normalized using total-count normalization to a target sum of 10,000 transcripts per cell, followed by log_2_-transformation. The data were then scaled to unit variance and zero mean for each gene.

##### Malignant and non-malignant cell discrimination

Cells were clustered using Louvain’s algorithm with a resolution of 0.5 ^99^. We performed differential expression analysis across clusters and used known tumor marker genes to classify cells as malignant or non-malignant.

##### Optimal Transport for annotation transfer between RNA-seq and Visium HD

Optimal Transport (OT) is a mathematical framework that originates from the problem of transferring one distribution of mass to another in the most cost-efficient manner. OT has become a foundational tool in fields such as economics, machine learning, and biology ^100–103^. In this work, we employed the Fused Gromov-Wasserstein distance coupled with spatial constraints ^104–108^, to learn a probability matrix 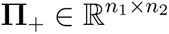 whose positive entry Π*_ij_* denotes the probability of transporting cell *i* to cell *j*, between two data modalities composed of *n*_1_ and *n*_2_ cells, respectively. Given a pairwise alignment matrix Π = [π*_i,j_*], mapping RNA-seq to Visium HD, we assigned to each non-malignant Visium HD cell, the cell type label of the most probable match from the RNA-seq data. The same procedure was repeated to assign meta-program labels to malignant Visium HD cells.

##### Malignant and non-malignant spatial interactions

To quantify spatial enrichment of malignant meta-programs and non-malignant cells, we developed a tumor neighborhood analysis framework based on Euclidean spatial distances between cell coordinates. For each non-malignant cell *i*, we first identified malignant neighbors within a fixed radius *r*. Let *N_i_*, denote the set of malignant neighbors of cell *i*, and let *T* be the set of malignant subtypes. For each subtype *t* ∈ *T*, we computed the number of neighbors of subtype *t* as 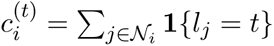, where *l_j_* is the subtype label of neighbor *j*, and **1**{·} is the indicator function. Aggregating these counts across non-malignant cells resulted in a tumor neighbor count matrix stratified by subtype. To assess statistical significance, we conducted permutation testing by shuffling the malignant subtype labels uniformly at random *N* = 100 times while preserving cell positions. For each permutation *n*, we recomputed neighbor counts 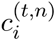 and calculated the mean 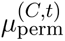 and standard deviation 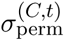 of neighbor counts for each non-malignant cell type *C*. The observed mean for cell type *C* and tumor subtype *t*, denoted 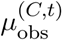, was then compared to the null distribution using a Z-score:

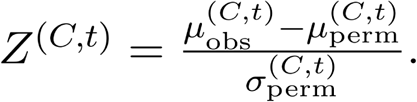

##### Peripheral Localization Index (PLI)

To quantify the spatial localization of malignant subtypes within the tissue architecture, we computed a Peripheral Localization Index (PLI), measuring the tendency of a cell cluster to localize near the tissue boundary. In detail, let 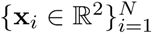 be the spatial coordinates of all cells, and let *C* ⊆ {1, …, *N*} denote the indices of cells in a target cluster. The boundary of the tissue is defined by the convex hull *H* of all cell coordinates. Let 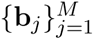 be the coordinates of the convex hull vertices. We first computed the average distance to the boundary as:

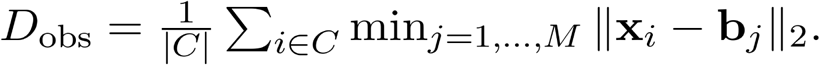

To construct a null distribution, we generated *K* = 500 random samples *R*^(1)^, …, *R*^(*k*)^, each containing |*C*| cells drawn uniformly without replacement from the full population. For each sample, we then computed:

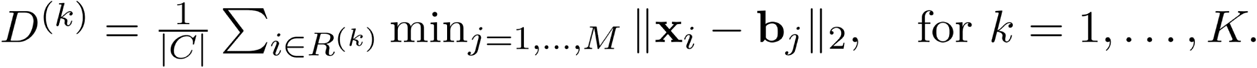

The mean and standard deviation of the null distribution are then:

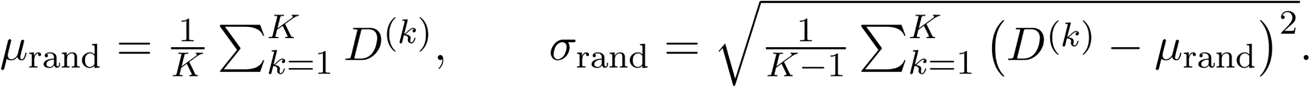

The Peripheral Localization Index (PLI) for a target cluster is finally defined as:

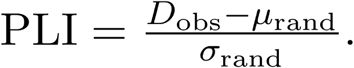

A negative PLI indicates that the cluster is significantly closer to the tissue boundary than expected by chance (i.e., peripheral enrichment), while a positive PLI suggests central localization.

##### Tumor State Proximity Score

To evaluate spatial clustering of phenotypically distinct tumor states, we computed a Tumor State Proximity Score (TSPS). For each tumor cell *i*, we defined its neighborhood *N_i_* as the set of cells located within a radius *r* in Euclidean space. Let *s_i_* ∈ *S* denote the tumor state label of cell *i*, and *s_j_* be the label of neighbor *j* ∈ *N_i_*. The per-cell TSPS is defined as:

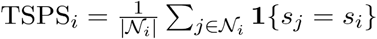

where **1**{·} is the indicator function. For each tumor state *s* ∈ *S*, we computed the average TSPS across all cells labeled with that state: 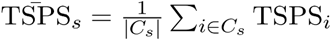, where *C_i_* = {*i* : *s_i_* = *s*} is the set of all cells assigned to state *s*.

##### Tumor Region Identification

For each sample, we delineated the tumor region directly from malignant cell coordinates. First, we estimated a continuous spatial intensity field using Gaussian kernel density estimation (KDE). The resulting density map was binarized with an adaptive threshold proportional to the mean density (τ·μ; τ = 0.5 by default). To regularize the mask, we removed small objects and holes and applied a binary closing followed by dilation. We then extracted the outer boundaries at the 0.5 isocontour level and converted them to polygons, discarding invalid or tiny shapes (area < 100). Finally, all valid polygons were merged to yield a single tumor contour.

##### Ecotypes Identification

We defined “ecotypes” as recurrent spatial composition patterns of cellular states across Visium HD slides. To identify globally consistent ecotypes, we defined a two-stages pipeline: *(i)* within-sample discovery of local ecotypes from spatial neighborhoods, and *(ii)* cross-sample consensus meta-clustering of local ecotype profiles to obtain study-wide, global ecotypes.

###### (i) Within-sample discovery of local ecotypes

For each Visium HD cell, we constructed a spatial k-nearest-neighbor (kNN) graph using Euclidean distance on the spatial coordinates (k = 50). Cell-type annotations were one-hot encoded, and for each cell we averaged the one-hot vectors of its kNN to obtain a local composition vector representing the proportions of cell types in its immediate neighborhood. To account for the compositional nature of these vectors, we applied a centered log-ratio (CLR) transformation. Cells were then clustered on the CLR-transformed features using K-means (n = 6 clusters per sample). For interpretability and downstream consensus analysis, we computed a mean neighborhood composition profile for each local ecotype by averaging the composition vectors of its member cells.

###### (ii) Cross-sample consensus ecotypes

To generate global ecotypes, we performed agglomerative hierarchical clustering on all per-sample ecotype profiles. Rather than fixing the number of consensus clusters, we selected a distance threshold by maximizing the Calinski–Harabasz (CH) index over a grid of candidate cuts. Specifically, we computed pairwise cosine distances, formed a grid of 25 thresholds at the 10th–90th percentiles of the empirical distance distribution, fit the hierarchy at each threshold, and retained the cut with the maximal CH on the original profile vectors. The selected threshold was used to obtain the final consensus ecotypes and their number K.

#### Transposable Elements (TEs) analyses

##### TE annotation

Repetitive DNA feature localizations (long interspersed nuclear elements, LINEs; short interspersed nuclear elements, SINEs; long terminal repeat elements, LTRs; and other categories such as DNA transposons and simple repeats) were retrieved from the UCSC RepeatMasker annotation (genome assembly GRCh38/hg38) using the AnnotationHub (v3.10.1) R package. The age of each TE instance was estimated based on sequence divergence, quantified as base mismatches per thousand (milliDiv) reported by RepeatMasker. Each milliDiv value was divided by the human genome substitution rate (2.2 × 10^−9^) to derive the final age ^109^

##### TE matrix generation

For both snRNA-seq and snATAC-seq datasets, MATES ^57^ was employed to estimate counts at the TE locus level from both unique and multi-mapping reads. Expression and accessibility matrices were subsequently generated by merging counts through summation, yielding a single matrix for each modality. Data were then normalized using size factors and log-transformed. The resulting matrices were normalized using size factors, log-transformed, and subsequently converted into Seurat object (v5.1.0) ^86^ for subsequent analysis and visualization.

##### Integration of modalities

TE-based RNA and ATAC modalities were integrated using the methods described above. Principal components (PCs) were selected based on the cellular subsets being analyzed. For the tumor microenvironment, RNA PCs 1-6 and ATAC PCs 1-10 were used. For malignant subsets, RNA PCs 1-9 and ATAC PCs 1-10 were considered for integration.

##### Differential expression and accessibility analyses

Differential expression and accessibility analyses between conditions were performed using the *scoreMarkers* function from the scran (v1.30.2) package. For each TE, rank-based statistics (i.e. *rank.AUC* and r*ank logFC.cohen*) were computed from both RNA and ATAC data to identify significant changes in expression and chromatin accessibility.

##### TE annotation on cCRE elements

TE loci overlapping MACS2-called peaks were further intersected with ENCODE cCREs and enhancer annotations from GeneHancer v4.4 ^110^. cCRE–gene associations were retrieved from the ENCODE database to link TE-associated regulatory elements with their potential target genes.

##### TE-Tumor Associated Antigens (TE-TAAs)

A computational workflow was developed to identify TE-TAAs. This workflow consists of the following steps:

1. Tumor-expressed and accessible TEs. Differentially expressed and accessible TEs in malignant cells compared to immune cells were identified. The top 1% of TEs, ranked by logFC Cohen metric for both modalities (as described in the differential analysis section), were selected for downstream analysis.
2. Detection of potential peptide-coding TEs. Nucleotide sequences of each tumor-associated TE were translated in all six reading frames using a custom R function. Only putative open reading frames (ORFs) of at least 10 amino acids were retained.
3. Filtering of false-positive TE-derived peptides. Candidate peptides were aligned with BLASTp ^111^ against methionine-initiated ORFs from the gEVE database ^112^. Only sequences with ≥95% identity and e-value < 0.01 were retained as high-confidence TE-derived peptides.
4. MHC-I binding prediction. Putative TE-derived antigens were analyzed using netMHCpan and prioritized based on both MHC-I binding affinity and recognition potential scores, following the procedure described in ^12^
5. High-confidence TE-TAAs. Finally, an antigen quality metric ^113^ integrating binding affinity and Neoantigen Recognition Potential (NRP), which estimates the probability that a presented neoantigen will be recognized by the TCR repertoire ^114^, was applied to define high-confidence TE-TAAs.

A total of 154 unique TE-derived antigens were ultimately identified, originating from 84 distinct TE loci.

To quantify the extent of malignant cells potentially undergoing immune elimination mediated by identified TE-TAAs, referred to as *immunoedited cells*, we calculated, for each MP, the percentage of cells that met the following criteria: (i) expression and chromatin accessibility at a TE locus giving rise to a TE-TAA (normalized counts > 0), and (ii) identification of the corresponding TE-TAA as immunogenic in the matched patient sample. This percentage was computed relative to the total number of cells belonging to the same MP within each patient. Changes in such percentage were evaluated across treatment time points, considering week 0 as baseline and combining post-treatment samples collected at week 4 and week 12.

#### Melanoma external cohort

##### Bulk RNA-seq melanoma cohorts under ICI therapies

Bulk transcriptomic data and clinical information were obtained for eight independent cohorts of melanoma patients treated with ICI. Six datasets ^5,115,116,117,118,119^ were acquired from the Tumor Immunotherapy Gene Expression Resource (TIGER) ^120^. Two additional datasets ^121,122^ were downloaded from cBioPortal ^123^ and from the Gene Expression Omnibus (GEO) under accession number GSE168204, respectively.

The composition of transcriptional metaprograms was inferred from the bulk RNA-seq expression matrix using the CIBERSORTx webserver ^124^. A custom signature matrix, generated from snRNA-seq data from this study, was used as the reference for deconvolution. The estimated tumor cell fractions were subsequently scaled based on tumor purity scores calculated with ESTIMATE (v1.0.13) ^125^.

##### scRNA-seq melanoma cohort under ICI therapies

A single-cell RNA-seq dataset of melanoma (EGA accession EGAS00001006488) ^16^ was analyzed. To improve data quality and reduce noise, samples containing fewer malignant cells than the 10th percentile were excluded, and genes expressed in less than 1% of cells were removed. Expression of MP signatures was quantified using the *AddModuleScore* function, and cells were assigned to the signature with the highest score, retaining only assignments with a score greater than 0.05.

#### Survival analysis

Samples were stratified into High- and Low- MP groups according to the median composition of each MP. Survival analysis was performed using the Kaplan–Meier method, with group differences assessed by the log-rank test. Cox proportional hazards regression models were fitted to estimate hazard ratios (HRs) and corresponding 95% confidence intervals. Results were visualized as forest plots. All statistical analyses were conducted in R using the survival package (v3.5-7) and the survminer package (v0.5.0).

#### Prognostic assessment

Prognostic value was evaluated in both the internal and external cohorts. For each sample, the relative proportions of metaprograms within the malignant compartment were calculated. Predictive performance for treatment response was estimated using leave-one-out cross-validation (LOOCV) with a generalized linear model implemented in caret (v6.0-94). Model accuracy was assessed by receiver operating characteristic (ROC) analysis, and the area under the curve (AUC) was computed using pROC (v1.18.5).

#### In-silico perturbation analysis

In-silico perturbation of selected TFs was performed using the SCENIC+ workflow ^67^. For each TF, simulated perturbation outputs were generated across five independent iterations. Within each iteration, the expression values of genes belonging to the *Neural crest-like* metaprogram signature were averaged and compared against the baseline expression in cells assigned to this transcriptional state. The mean difference across signature genes was then computed per iteration, yielding an overall measure of the perturbation effect for each TF.

#### Gene regulatory network inference in external scRNA-seq melanoma dataset

GRNs were inferred from the external scRNA-seq dataset ^16^, using pySCENIC (v0.12.1+8.gd2309fe) ^97^. To obtain a high-confidence network, the initial regulons were refined by applying a weight-based cutoff: for each TF, the 90th percentile of its target connection weights was used as a threshold, and only interactions exceeding this value were retained.

#### NFATC2 post-transcriptional activity

NFATC2 regulon activity was quantified on bulk RNA-seq datasets using the univariate linear model (ULM) framework implemented in decoupleR (v2.9.7) ^126^. The resulting activity score was subtracted from NFATC2 expression values to derive its post-transcriptional activity.

The predictive capacity of NFATC2 was assessed in pre- and post-ICI samples independently using LOOCV with a GLM as described above. Model performance was evaluated with ROC analysis and the corresponding AUC using pROC.

#### NFATC2 knockdown in melanoma cell lines

Gene expression data for NFATC2 knockdown and control melanoma cells were obtained from GEO (GSE101323) ^75^. Differential expression was performed with limma (v3.58.1) ^127^. Gene set enrichment analysis was carried out using Gene Ontology Biological Process, KEGG, Reactome, WikiPathways, and Hallmark collections (as described above). A single-sample Mann–Whitney–Wilcoxon Gene Set Test (MWW-GST) ^128^ was applied using the top 50 up-regulated genes for each MP.

## Acknowledgements

NIBIT Foundation Onlus; Fondazione AIRC under 5 per Mille 2018-ID.21073 project - P.I. Maio Michele, G.L. Anichini Andrea; National Center for Gene Therapy and Drugs based on RNA Technology (Project no. CN_00000041) European Union -Next Generation EU, Mission 4, Component 2, CUP B93D21010860004 Maio Michele, CUPE63C22000940007. Research reported in this publication was performed in part at the Cancer Modeling Shared Resource (CMSR; RRID: SCR02289), the OncoGenomics Shared resource (OGSR; RRID: SCR022502) and the Biostatistics and Bioinformatics Shared Resource (BBSR; RRID:SCR022890) of the Sylvester Comprehensive Cancer Center at the University of Miami Miller School of Medicine, which is supported by the National Cancer Institute Cancer Center Support Grant (CCSG) P30-CA240139.

## Supplementary Figure Legends

**Supplementary Figure 1.**
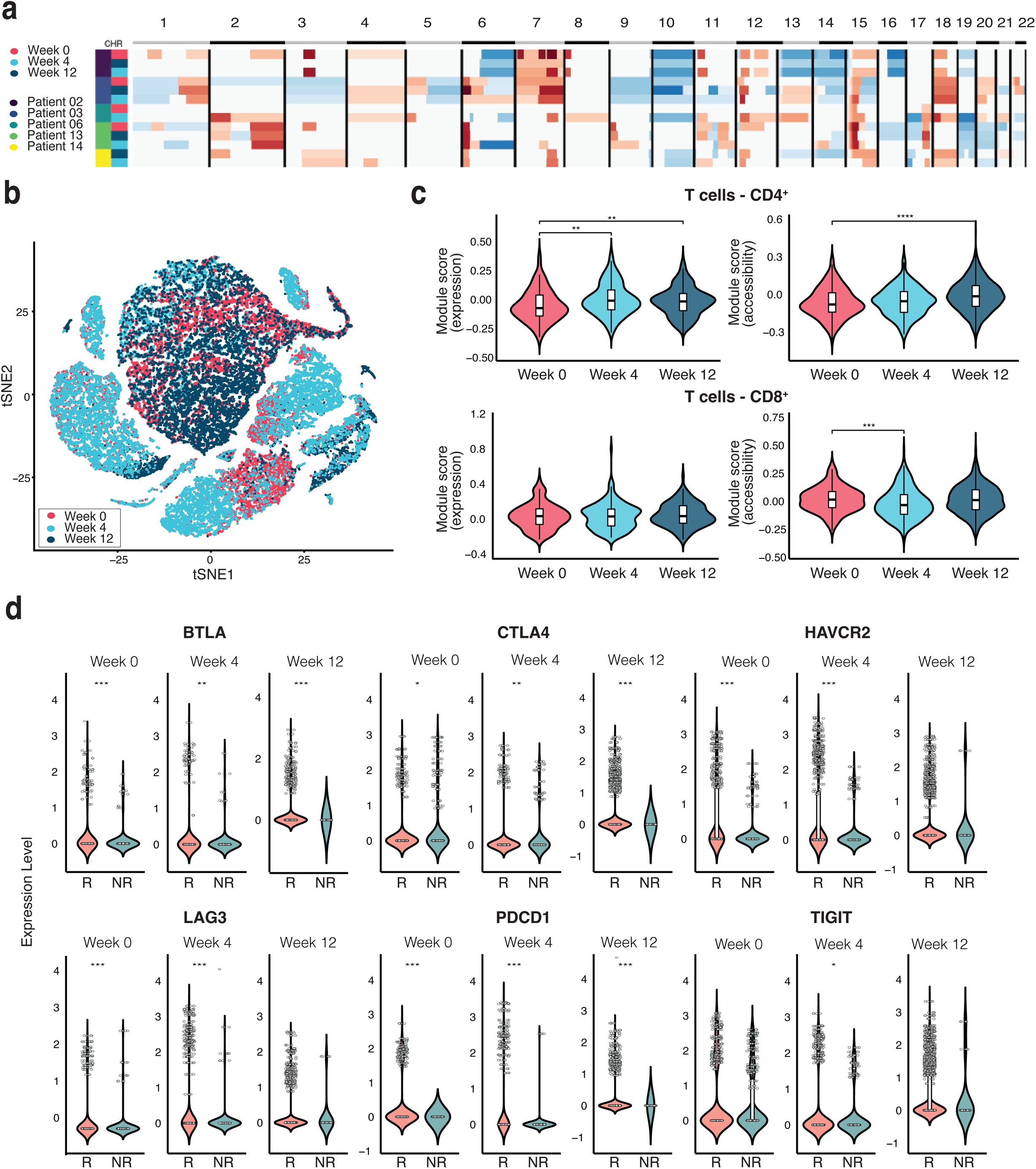
Genomic and immune landscapes of the NIBIT-M4 cohort, related to Figure 1. **a.** Consensus heatmap of Copy Number Alterations (CNAs) in malignant cells inferred with SCEVAN ^18^; patient IDs and sample time points are shown on the left track. CNAs are color-scaled from –2 (deep deletion, blue) to +2 (high amplification, red). **b.** t-SNE visualization of co-embedding of snRNA and snATAC modalities colored by time-points. **c.** Violin plots of Reactome PD1-signaling module scores from expression (left) and gene accessibility (right) in CD4^+^ (top) and CD8^+^ (bottom) cells across time-points. Statistical significance was assessed using Wilcoxon rank-sum test; ** *p* ≤ 0.01, *** *p* ≤ 0.001, **** *p* ≤ 0.0001. **d.** Expression of immune checkpoint genes in R and NR patients across time-points Statistical significance was assessed using Wilcoxon rank-sum test; * *p* ≤ 0.05, ** *p* ≤ 0.01, *** *p* ≤ 0.001.

**Supplementary Figure 2.**
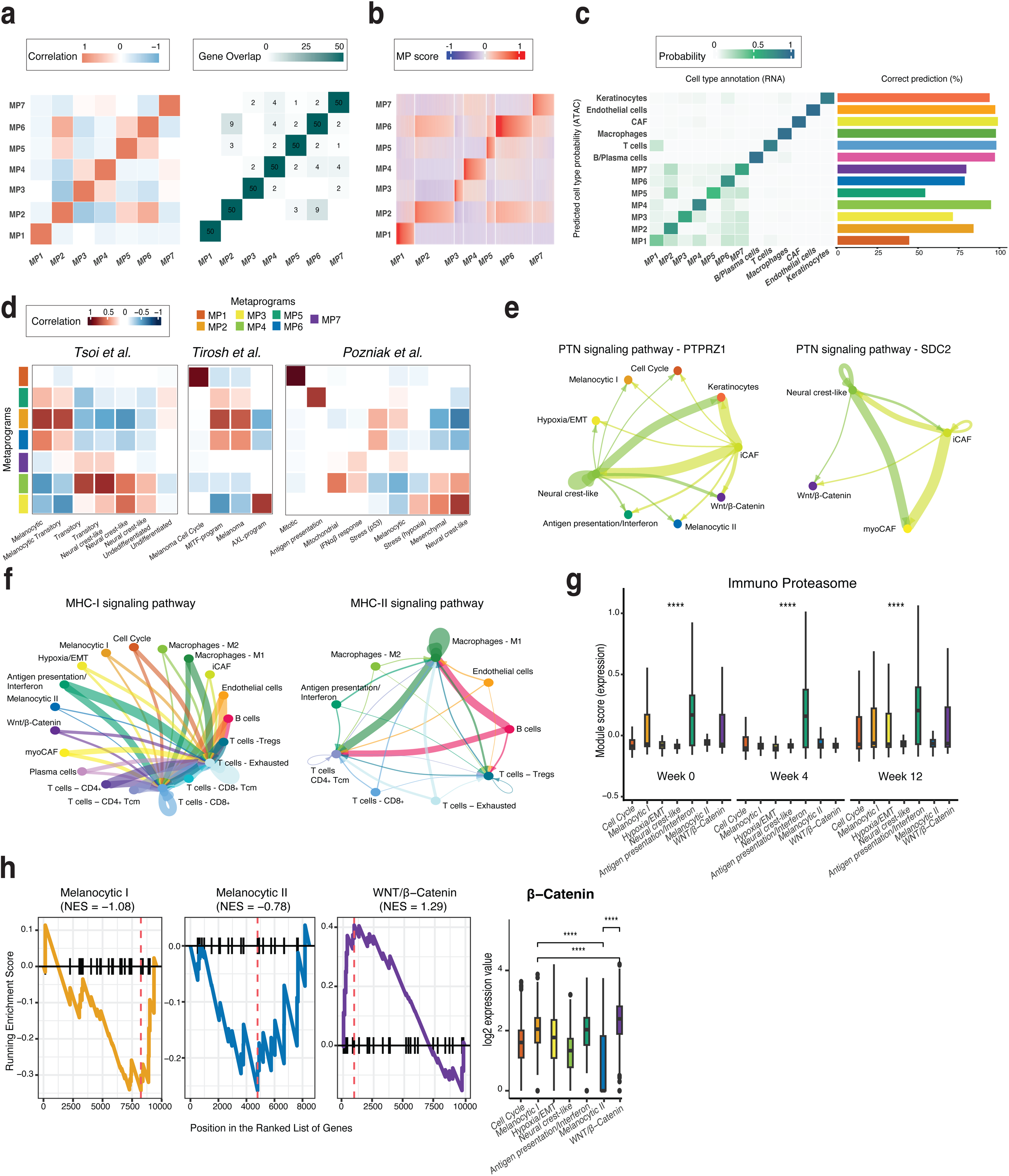
Functional dissection of metaprograms, related to Figure 2. **a.** Heatmaps showing Pearson correlations between each MP score and all other MP scores (left), and 50-NMF gene overlap (right) between MPs. **b.** Heatmap of MP scores across cells. Rows represent MPs, columns represent cells. Cells were aggregated by MPs and ordered according to their scores. **c.** Heatmap of average percentage score of cells from a given expression-defined state (columns) assigned to different cell states (rows) based on gene accessibility profiles, with barplot of prediction accuracy. **d.** Heatmaps showing Pearson correlations between MP scores with previously published melanoma signatures ^4,14,16^. **e.** CellChat aggregated networks of selected pathways in NR patients; line width indicates interaction probability. **f**. Same analysis as in (**e**), shown for R patients. **g.** Boxplots of immunoproteasome (*PSMB8, PSMB9, PSMB10*) module scores across MPs at different time-points. Statistical significance was assessed using Wilcoxon rank-sum tests; **** *p* ≤ 0.0001. **h.** Wnt/β-catenin program characterization with enrichment plots of HALLMARK_WNT_BETA_CATENIN_SIGNALING pathway in Melanocytic I, Melanocytic II and Wnt/β-catenin MPs, and β-catenin gene expression (log2 normalized counts). Statistical significance was assessed using Wilcoxon rank-sum tests; **** *p* ≤ 0.0001

**Supplementary Figure 3.**
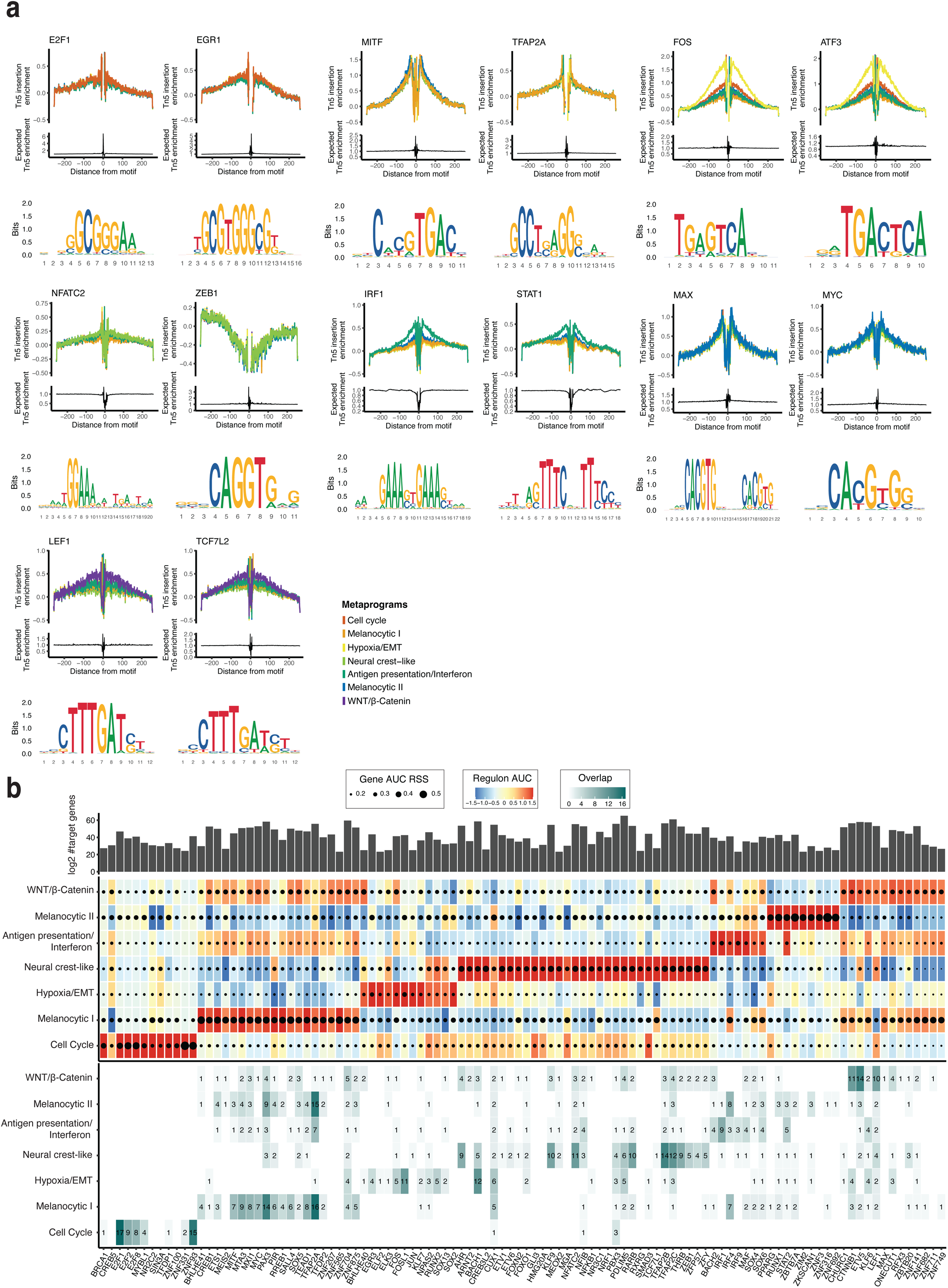
Transcription factor activity across metaprograms, related to Figure 2. **a.** Footprinting analysis of representative TFs per MP showing chromatin accessibility patterns (top) together with sequence logos of their binding motifs (bottom). **b.** TF eRegulon profiles across MPs. Top: bar plot of eRegulon size represented as log2 number of target genes. Middle: heatmap/dot plot displaying TF expression (AUCell score, color scale) and eRegulon cell-type specificity measured by the regulon specificity score (RSS; point size). Bottom: overlap analysis between each TF eRegulon and the top 50 signature genes of MPs, shown per TF column.

**Supplementary Figure 4.**
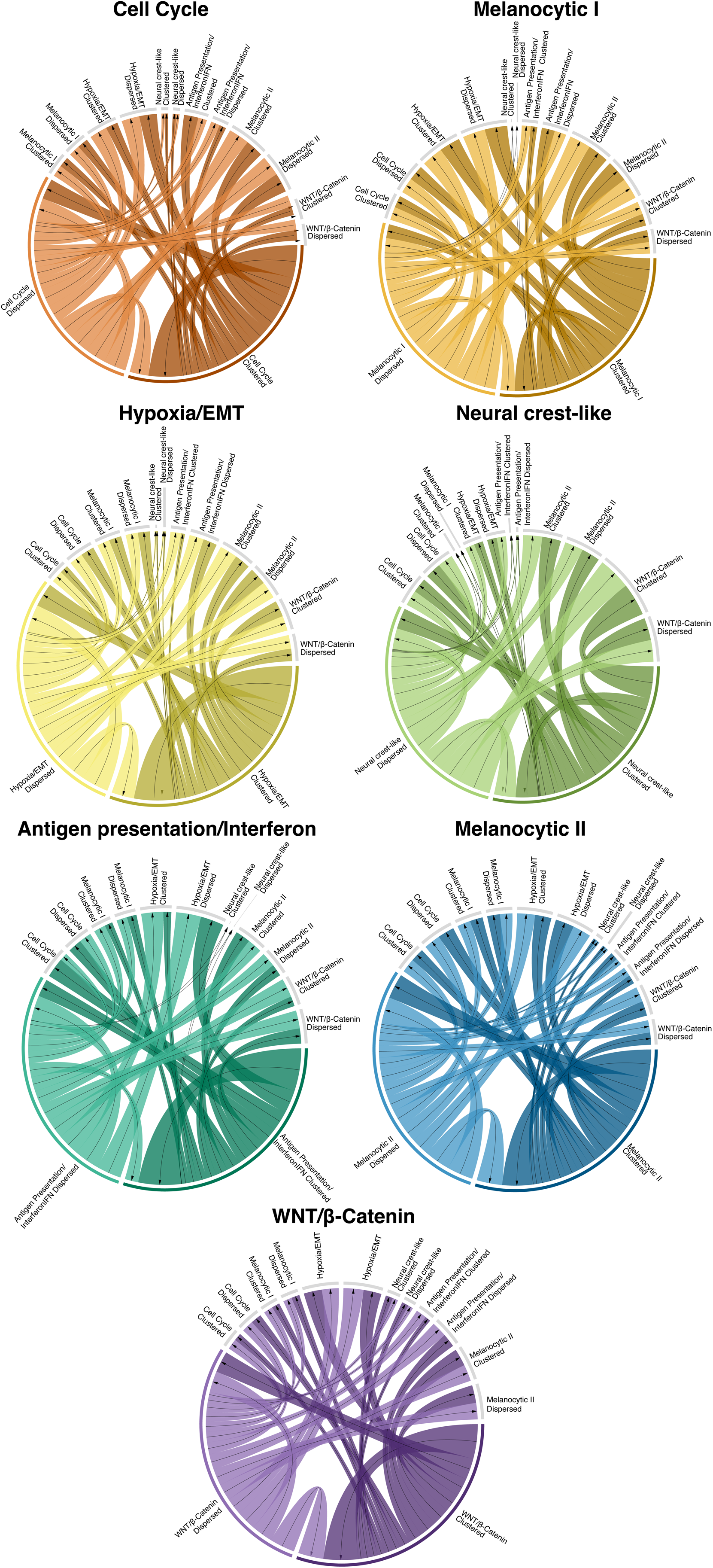
Transition probabilities for the malignant cell states conditioned on their spatial configuration. For all metaprograms, the clustered cells showed a higher probability to transition into the same state and retain their spatial organization compared to the dispersed states. The strongest homotypic transitions were observed in the Antigen presentation/Interferon and Hypoxia/EMT *states*.

**Supplementary Figure 5.**
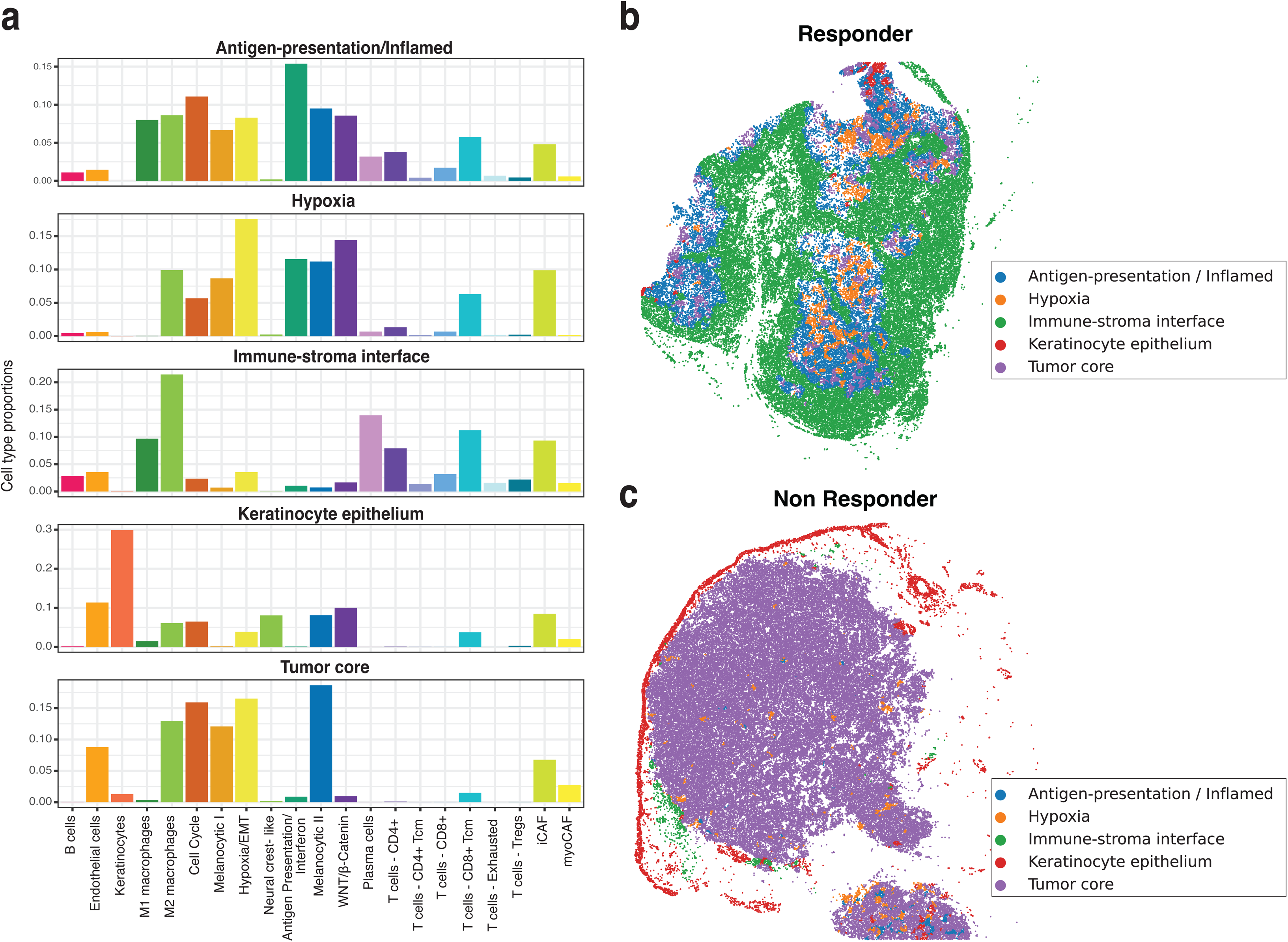
Unsupervised ecotypes from Visium HD samples. **a.** Proportions of cell types in each ecotype. The ecotype composition was used to label them as Antigen-presentation/Inflamed, Hypoxia, Immune–stroma interface, Keratinocyte epithelium, and Tumor core. **b.** Spatial distribution of the identified ecotypes in a responder and non-responders sample **c.**, respectively.

**Supplementary Figure 6.**
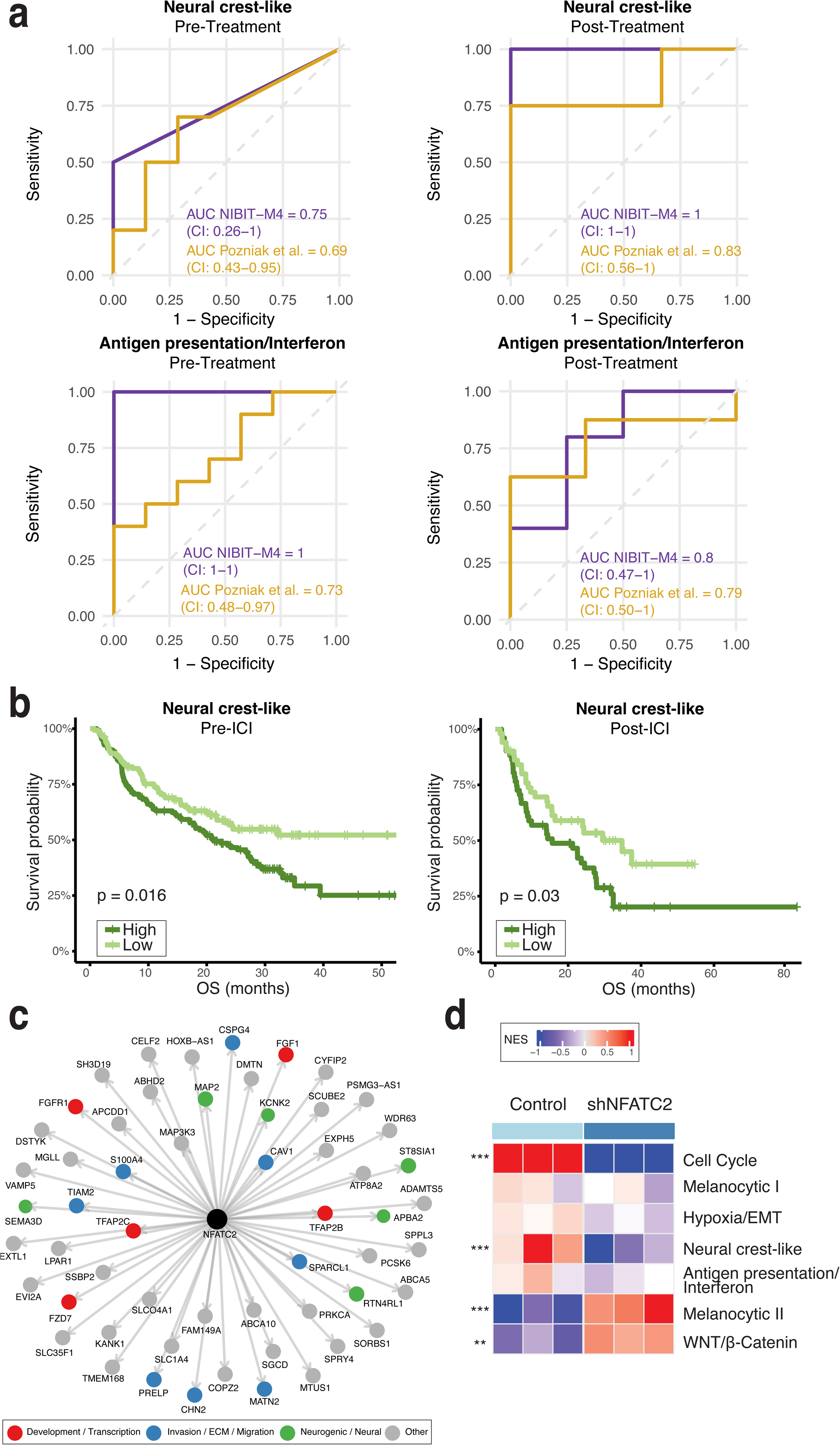
Prognostic significance of metaprograms and NFATC2 functional regulation, related to Figure 6. **a.** Receiver operating characteristic (ROC) curves of response prediction for the fraction of *Neural crest-like* (top) and *Antigen presentation/Interferon* (bottom) cells among malignant cells in the NIBIT-M4 cohort (violet) and an external cohort ^16^ (goldenrod), shown pre-treatment (left) and post-treatment (right). Areas under the ROC curve (AUCs) are reported with 95% confidence intervals (CIs). **b.** Kaplan–Meier curves of overall survival (OS) in bulk melanoma cohorts, shown separately for pre-ICI (left) and post-ICI (right). Patients were divided into High (dark green) and Low (light green) groups based on the median fraction of Neural crest-like cells. Statistical significance was assessed using the log-rank test. **c.** NFATC2 regulon network showing gene targets as nodes, colored by major functional category. **d.** Heatmap of single-sample normalized enrichment scores (NES) for the top 50 DEGs per MP in control and shRNA NFATC2 conditions. NES values are displayed on a –1 to +1 color scale (blue to red). Statistical significance was calculated using combined *p*-values across replicates with Fisher’s method; ** *p* ≤ 0.01, *** *p* ≤ 0.001.

